# Rational programming of history-dependent logic in cellular populations

**DOI:** 10.1101/617209

**Authors:** Ana Zúñiga, Sarah Guiziou, Pauline Mayonove, Zachary Ben Meriem, Miguel Camacho, Violaine Moreau, Luca Ciandrini, Pascal Hersen, Jerome Bonnet

## Abstract

Genetic programs operating in an history-dependent fashion are ubiquitous in nature and govern sophisticated processes such as development and differentiation. The ability to systematically and predictably encode such programs would advance the engineering of synthetic organisms and ecosystems with rich signal processing abilities. Here we implement robust, scalable history-dependent programs by distributing the computational labor across a cellular population. Our design is based on recombinase-driven DNA scaffolds expressing different genes according to the order of occurrence of inputs. These multicellular computing systems are highly modular and any program can be built by differential composition of strains containing well-characterized logic scaffolds. We developed an automated workflow that researchers can use to streamline program design and optimization. We anticipate that the history-dependent programs presented here will support many applications using cellular populations for material engineering, biomanufacturing and healthcare.

**One Sentence Summary:** Systematic and automated frameworks for implementing robust history-dependent genetic programs in cellular populations.

## Main Text

Living organisms execute sophisticated functions and generate complex structures at all scales (*1*–*3*). These rich, robust, and reliable behaviors are often achieved through division of labor, as seen in bacterial biofilms, multicellular organisms, or insect colonies (*4*–*7*). Synthetic biologists have reproduced complex multicellular behaviors such as social interactions (*8*–*10*), oscillations (*11*, *12*), or spatial patterning (*13*–*18*).

Programming living systems requires signal processing devices integrating environmental signals and producing a response according to a pre-defined logic algorithm (*19*). Logic devices executing Boolean functions in living cells have been engineered using a wide range of biochemical mechanisms (*20*, *21*). Taking inspiration from the division of labor found in natural system, Boolean logic was also achieved at the multicellular level by distributing the computational labor across various strains (*22*–*25*).

A key feature of biological systems is to respond to signals according to the order in which they are received. History-dependent response is ubiquitous in biology and is observed from animal behavior down to the heart of fundamental processes like cell division (checkpoints), differentiation (cell-fate commitment) and development (*26*–*28*). History-dependent behavior was also proposed to be important for microbial survival strategies and to provide a fitness advantage in the evolutionary competition (*29*, *30*). Similarly, human-engineered devices rely on sequential logic circuits to execute complex programs (*31*).

The ability to implement history-dependent genetic programs within living cells has many practical implications from a research and engineering perspective. Such synthetic programs could be used as temporal and spatial trackers for decoding biological processes such as development. Furthermore, living organisms could be programmed to exhibit sophisticated behaviors not found in nature. History-dependent circuits could be applied in biomanufacturing to mediate sequential, substrate triggered activation of components of a metabolic pathway. Engineered bacterial therapeutics could also be embedded with history-dependent programs that would improve their safety and specificity. For instance, bacterial cancer therapeutics could be activated only when they have encountered certain body location in a particular order and reached a high density inside the tumor, possibly in combination with external therapeutic triggers (*32*–*35*). Furthermore, history-dependent programs would be instrumental to implement morphogenetic programs for engineering synthetic tissues (*36*) and living functional materials which require order-specific, sequential assembly (*37*–*40*).

Scientists have started to explore the engineering of history-dependent biological programs, using different molecular memory devices to store the order of occurrence of signals. For instance, a mutual inhibition memory device (*41*, *42*) was used as the basis for building a Push-on/ Push-off switch (*43*), a device toggling between two states in response to the same signal, or to engineer a circuit producing a response comparable to Pavlovian behavior in *Escherichia coli* (*44*). RNA devices were engineered to produce a response after a certain number of stimuli (*45*), and logic circuits based on transcriptional set-reset latches were used to implement synthetic cellular checkpoint control (*46*).

Among the different biochemical mechanisms to implement history-dependent behavior, memory devices using DNA recombination quickly emerged as a tool of choice. Recombinase memory devices use permanent inversion or excision of DNA sequences via site-specific recombinases (*47*, *48*). The state of the system can be encoded both by gene expression state and within the target DNA sequence. Contrarily to feedback-based systems, recombinase memory devices do not require constant protein production from the cell to hold state, reducing metabolic burden and increasing evolutionary stability of the engineered systems (*49*, *50*). Recombinases can encode complex Boolean logic within living cells using reduced, single layer architectures (*51*, *52*). By interleaving target sites for different recombinases, recombination reactions can be made dependent on each other, and the system can transition through different DNA states depending on which recombination reaction occurs first (*50*, *53*). Using this concept, researchers started to implement genetic devices tracking the order of occurrence of signals, as well as history-dependent gene expression programs (*50*, *54*, *55*). These history-dependent logic devices operate at the single-cell level, and the most advanced versions use multiple mutant recombination sites recognized by the same enzyme (*55*). However, current programs are designed in an *ad-hoc* manner, limiting their predictability and complicating their implementation. In addition, the concatenation of almost identical sequences (50bp mutant sites differ only by their central dinucleotides) might impact the scalability of these designs while creating DNA synthesis and evolutionary stability problems.

On the other hand, distributing the computational labor between different cells has proven to be a powerful approach to systematize circuit design, reuse biological components, and obtain predictable behavior from engineered biological systems. A particular advantage of distributed multicellular computation (DMC) is the high-number of logic functions achievable from a reduced collection of well characterized genetic devices (*22*, *23*, *25*). DMC could thus offer an attractive alternative to systematically implement history-dependent cellular programs in a predictable manner.

Here we provide a systematic framework for reliably implementing history-dependent programs within a cellular population by distributing the computation between different strains. Our design is based on modular scaffolds in which gene expression in particular states is linked to a specific scaffold recombination state. We demonstrate reliable execution of 2- and 3-input programs by engineered multicellular systems and show how circuits can be minimized by combining history-dependent and Boolean logic devices. We provide a web-based, automated circuit design tool along with a device optimization framework which are freely available to the community. We anticipate that the multicellular history-dependent logic presented here will support many applications in material engineering, biomanufacturing and healthcare.

### A modular scaffold design for history-dependent programs

History-dependent gene expression programs can be represented as a lineage tree (*56*) in which each branch, or lineage, corresponds to a specific order of occurrence of the inputs with the number of lineages being equal to N!, where N is the number of inputs (Fig. 1A). For instance, for 2-input programs, 2 lineages exist, while for 3-input programs, 6 lineages exist. We implemented history-dependent programs by using site-specific recombinases that perform DNA inversion and excision events (Fig. 1B); we focused on serine integrases which operate in several species and for which many orthogonal enzymes have been characterized and used to build genetic circuits (*25*, *51*, *52*, *55*, *57*).

**Figure 1.**
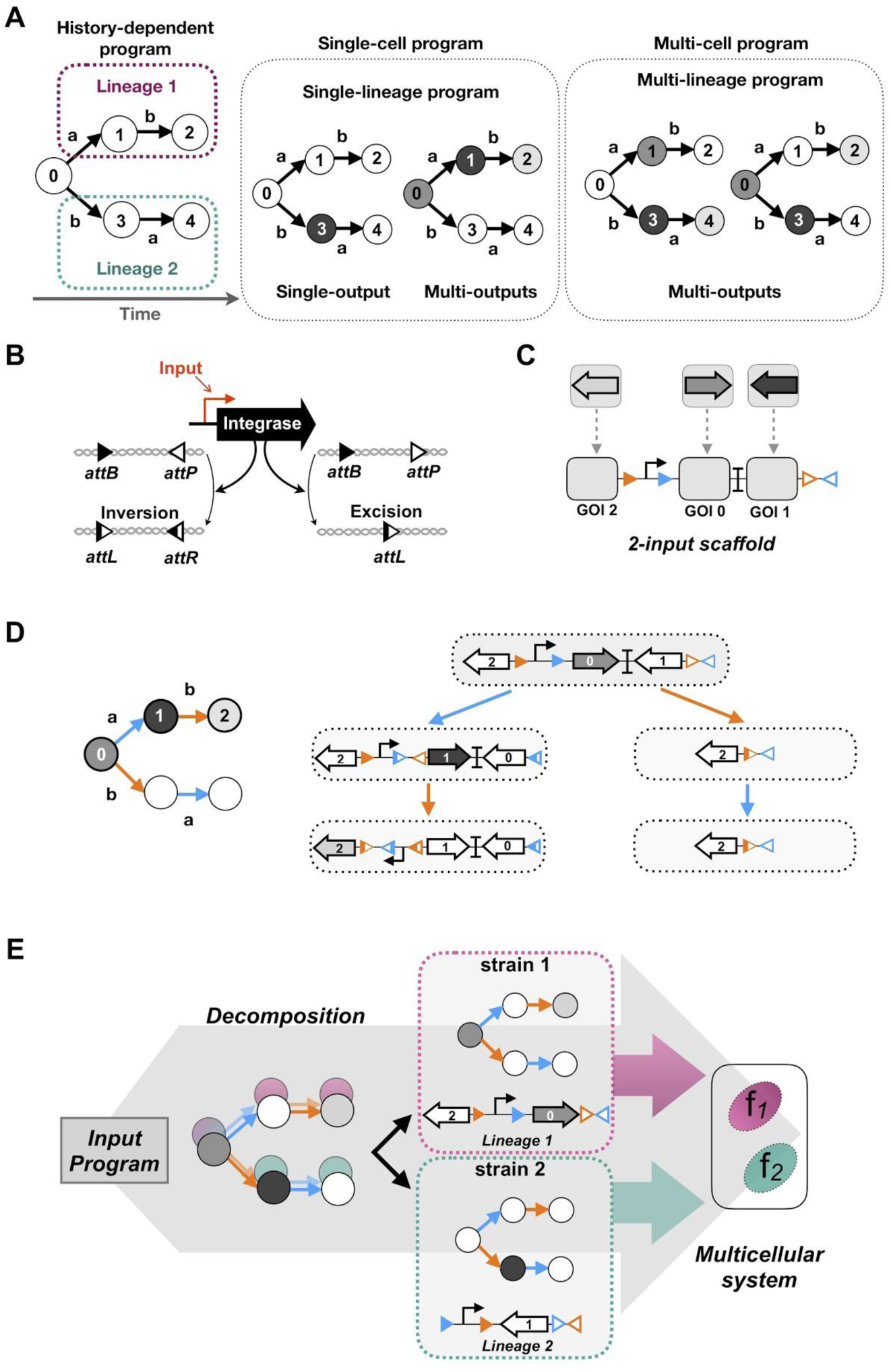
Design of a modular scaffold for 2-input history-dependent multicellular programs. (**A**) Lineage tree representing a history-dependent program. Letters represent the presence of the two inputs (a and b) and numbers on nodes represent states of the system associated with the order-of-occurrence of the inputs. For two inputs programs, five states are possible. For implementation of single-lineage programs one strain is required. For multi-lineage program, multiple strain are required. (**B**)Integrase-mediated excision or inversion. When integrase sites are in the opposite orientation (left panel), the DNA sequence flanked by the sites is inverted. If integrase sites are in the same orientation (right panel), the DNA sequence flanked by the sites is excised. (**C**) 2-input history-dependent scaffold. Integrase sites are positioned to permit expression of an output gene (black arrows) or not (empty gray squares) in the corresponding lineage. The programs are implementable based on this 2-input scaffold by inserting genes corresponding to the ON state in the corresponding positions. (**D**) Lineage tree representing the scaffold history-dependent program. For each state of the lineage (a then b) a different gene is expressed. On the right panel, the gene at the GOI position 0 is expressed only when no input is present. If input *a* is present first an inversion occurs, and gene 1 is expressed. If input *b* is present first and excision occurs, no gene is expressed (nor will be expressed) as the promoter is excised. If input *b* follows input *a*, gene at the position 2 is expressed. The scaffold show 4 different DNA states. (**E**) A multi-lineage program is decomposed into sub-programs, each sub-program corresponding to a different lineage (a then b; b then a). Each sub-program is implemented in a different cellular computation unit (CCU) as a DNA device (*f*1, *f*2). The composition of the CCUs into a multicellular system permits implementation of the full program.

We designed a modular scaffold capable of executing all possible 2-input history-dependent gene expression programs that can occur within a single lineage (Fig. 1C). The modular scaffold contains insertion sites or “slots” in which genes of interest (GOI) can be inserted and expressed in each particular state of the lineage tree (Fig. 1C). Each input is assigned to an integrase that controls recombination of specific portions of the scaffold. Using this scheme, any possible combination of gene expression states within a particular lineage can be achieved by simply inserting the desired gene at the corresponding positions, and switching the identity of recombination sites according to the desired lineage (Fig. 1D).

Depending on the identity of the different GOIs, the scaffold can express a single or multiple genes in the different output states. Programs requiring gene expression in different lineages are decomposed in sub-programs that will be executed by different strains, or cellular computation units (CCUs), each CCU executing one lineage. The full program is executed by a multicellular system obtained by mixing CCUs in equal proportions (Fig. 1E). For a given number of inputs, the maximum number of CCU needed is equal to the number of lineages (N! for N inputs) (Fig. S1). However, most functions are implementable with fewer than the maximum number of strains, as the number of CCUs depends on the number of lineages in which gene expression is required.

### Implementing single-lineage programs

We first implemented single-lineage 2-input programs. Such programs could be used for instance to track if a cell has entered a specific differentiation pathway. In order to streamline the engineering process, we took advantage of the similarity of recombinase systems with state machines and designed an optimization workflow called OSIRiS (<u>O</u>ptimization via <u>S</u>ynthesis of Intermediate <u>R</u>ecombination <u>S</u>tates). Because each state actually corresponds to a physically different DNA molecule which sequence can be fully predicted, OSIRiS enables the different recombination intermediates to be generated *in silico*, synthesized, and characterized (Fig. S2). OSIRiS therefore decouples scaffold optimization from integrase mediated switching, simplify and shorten the characterization process, and allows for rapid iteration cycles. The OSIRiS script, written in python, automatically generates the intermediate sequences of any integrase-based scaffold, and is available on github (*58*).

We then characterized the history-dependent response of nine single-lineage programs representing various combinations of ON states, lineages and output types (Fig. 2). Single-lineage programs with single-output permits to identify particular states the cell entered, while those with multi-outputs can enable precise, unambiguous discrimination between different states. For one lineage (lineage 2: b then a), we tested all possible history-dependent programs producing a single type of output (GFP). For lineage 1 (a then b), we tested programs using a different fluorescent reporter for each state. We used a dual-controller plasmid (*51*) in which Bxb1 integrase is under the control of the P_Tet_ promoter responding to anhydrotetracycline (aTc, input a) and Tp901-1 integrase is under the control of the P*_BAD_* promoter responding to arabinose (Ara, input b) (Fig. 2). For programs expressing GFP in multiple output states, we used different GFP variants with reduced sequence homology (*59*) to avoid recombination and replication slippage issues (*19*) (see supplementary file). We operated the system in fundamental mode, *i.e.* inputs cannot occur simultaneously, but sequentially (*31*). All nine single-lineage programs behaved as expected. The scaffold was capable of driving expression of various fluorescent reporters in different DNA states and in both lineages. All devices had at least a 10 fold change in fluorescence intensity between the OFF and ON state, with a max fold change over 250 (Fig. S3 and S4). We observed, as expected, variations in fold changes depending on which GFP variant was used. We also measured the percentage of switched cells, and found a minimal of 90% switching efficiency for states supposed to express a fluorescent protein. We also observed spontaneous switching in some states (mostly in S0) affecting a maximum of ∼12% of the population (Fig. S4).

**Figure 2.**
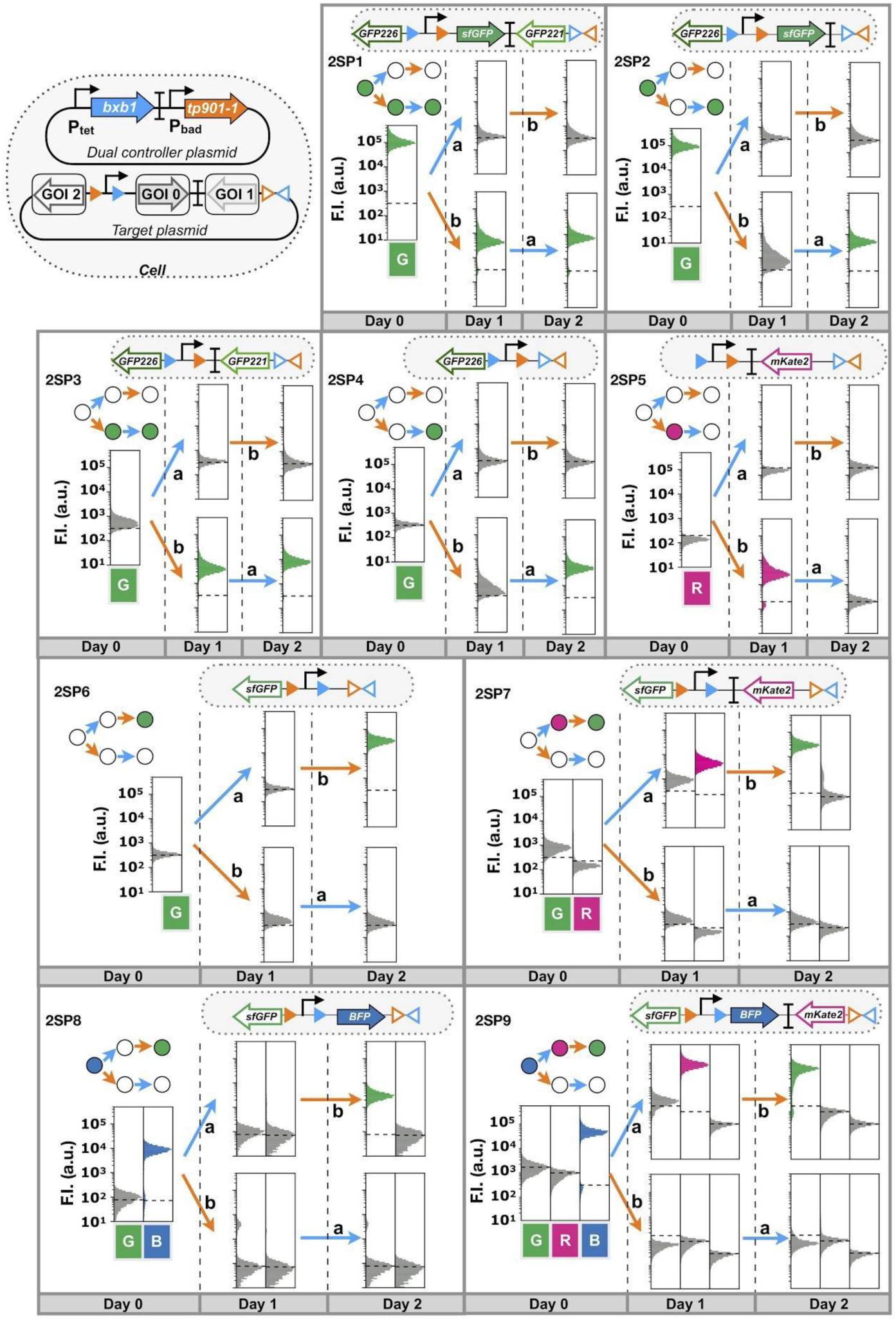
Characterization of 2-input single-lineage programs (2SP1-2SP9). Upper left: genetic design of the two plasmid used to characterize each program. The dual controller plasmid regulate the gene expression of the Bxb1 and Tp901-1 integrases. The target plasmid corresponds to the 2-input history-dependent scaffold. Nine single-lineage history-dependent programs exhibiting lineage specific, or state-specific gene expression with single or multiple genetic outputs were implemented and characterized. Bxb1 and Tp901-1 are induced by aTc (input a) and by arabinose (input b) respectively. The lineage trees for each program and its corresponding genetic DNA device are represented. Cells transformed with both plasmids were induced sequentially twice for 16 hours. Each histogram shows the expression of fluorescent reporters expressed at different induction stages. A representative example is depicted here. Fold change measurements can be found in Figure S4.

One particular program expresses a different fluorescent reporter gene in each input state of one lineage (Fig. 2, Fig S3A, program 2SP9). We obtained the expected phenotype for each of the four possible DNA states, BFP in state 0, RFP in state 1, GFP in state 2, and no expression in state 3 or 4 (Fig. S3B). Additionally, we analyzed switching kinetics of the 2SP9 program by time-lapse microscopy, and observed a clear transition of states from blue to red to green fluorescence. We noted that 3 h were sufficient to observe 100% switching in the lineage states from S0 to S1 in strains growing continuously in a mother machine microfluidics device (Fig. S5 and Supplementary Movie 1).

### History-dependent logic in multicellular systems

We then implemented 2-input history-dependent programs producing outputs in the 2 different lineages, therefore requiring the assembly of a multicellular system. As a demonstration, we built two 2-Input Multicellular Programs, 2MP1 and 2MP2, using four different CCUs with two strains per program (Figure 3). We co-cultivated the two strains for three days and subjected them to the different sequences of inputs.

**Figure 3.**
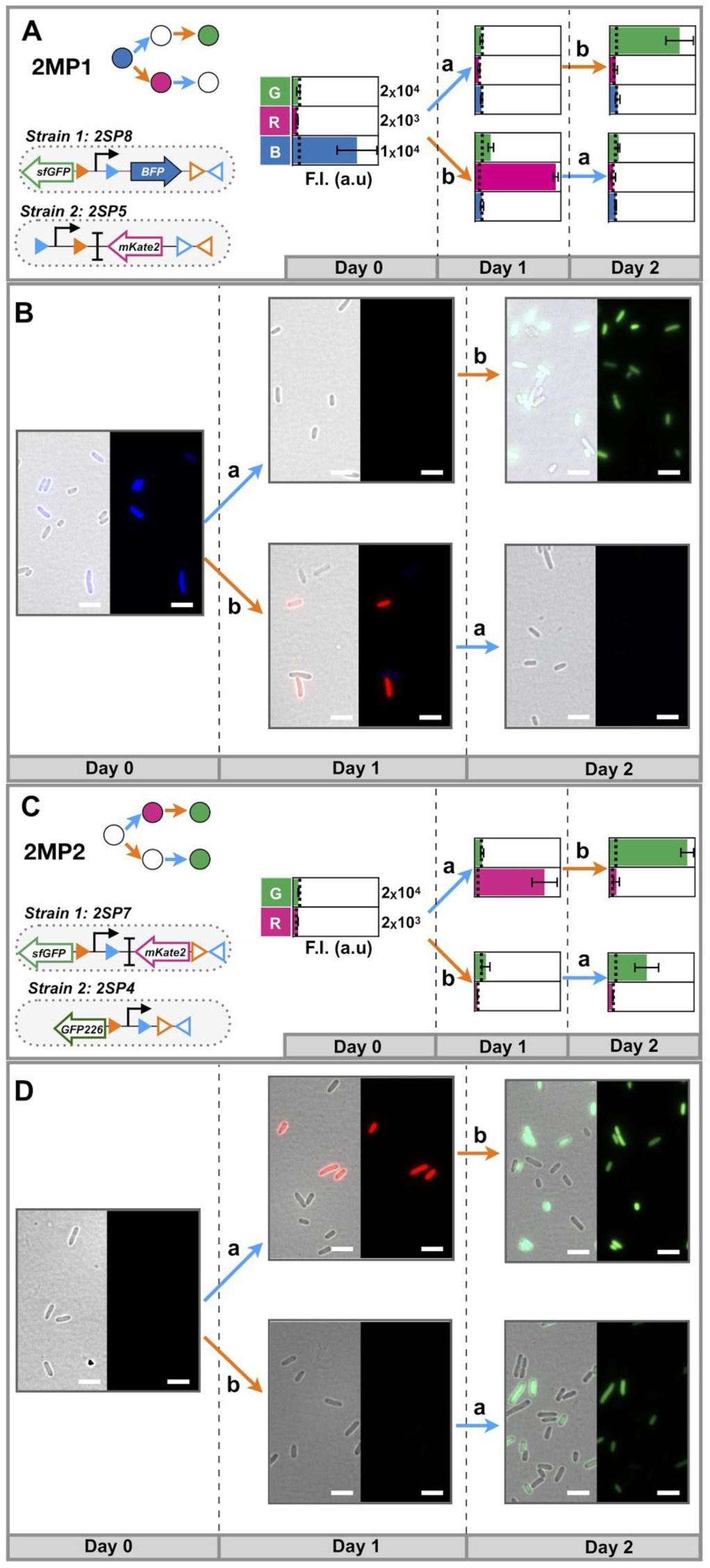
Characterization of 2-input multicellular programs. Design and characterization of 2-input Multicellular Program (2MP) 1 and 2, by plate reader (**A**, **C**) and microscopy (**B**, **D**), respectively. Both multicellular programs were implemented using two different single-lineage strains. The inputs are represented by letters, *a* for aTc inducing Bxb1 Integrase and *b* for arabinose inducing Tp901-1 integrase. To implement each program the CCUs were mixed in similar proportions and grown for 16 hours with each inducer. The bar graph corresponds to the fluorescence intensity (F.I) in arbitrary units (a.u) for each fluorescent channel (GFP, RFP and BFP), plotted in linear scale. The error bars correspond to the standard deviation for two different experiments. The dotted line indicates the negative autofluorescence from control strain. The microscopy images correspond to the merged images of the GFP, RFP, and BFP channels with bright-field (left) and without (right). Bars, 10 μm.

We measured the bulk fluorescence intensity of the cellular population in each state using a plate-reader, and confirmed that programs behaved as expected, with measurable, successive expression of different genes according to the order of inputs (Fig. 3A and 3C). Microscopy analysis showed, as expected, mixed gene expression states (ON and OFF) between the two strains (Fig. 3B and D). Flow cytometry analysis of both multicellular systems showed an equivalent percentage (∼50%) of cells in each subpopulation during three days of sequential induction, confirming the stability of the system and the lack of competition effects between the two strains (Fig. S6). The decrease in fold change resulting from strain dilution within the multicellular system could be predicted from individual strain measurements (Fig. S7).

We then sought to scale-up the system, and designed a 3-input scaffold capable of expressing a different gene in all states of a single 3-input lineage (Fig 4A-C). We added a third integrase, Integrase 5 (Int5) (*57*) to the controller plasmid, under the control of the pBEN promoter responding to benzoate (input c) (Fig. 4D, Fig. S8). We used OSIRiS to characterize and optimize the 3-input scaffold and its five recombination intermediate states (Fig. S9 and S10).

**Figure 4.**
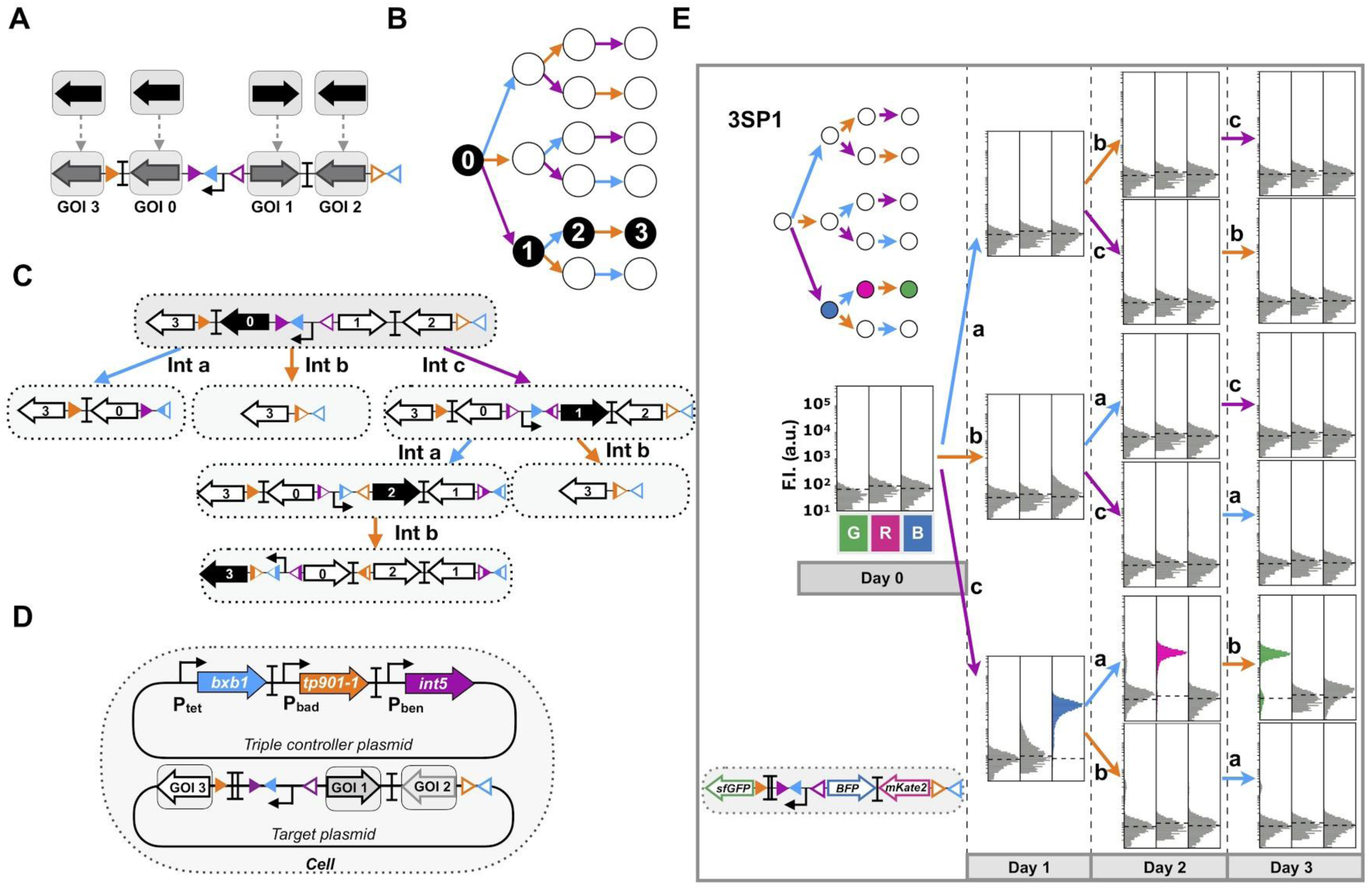
Design and characterization of a modular scaffold for 3-input history dependent programs. 3-input history-dependent scaffold (**A**) and its representative lineage tree (**B**). Integrase sites are positioned to permit expression of an output gene in the corresponding lineage. For each state of the desired lineage, a different gene is expressed, and a gene is also expressed when no input is present. The four rows of the lineage tree correspond to different numbers of inputs that have occurred sequentially (from 0 to 3 inputs) and the 6 lineages correspond to different order-of-occurrences of inputs (example: a-b-c for lineage 1 and b-a-c for lineage 3). (**C**) DNA states of the scaffold. The gene at the GOI position 0 is expressed only when no input is present. The scaffold show 5 different DNA states. (**D**) Genetic design of the two plasmids used to characterize 3-input programs. The triple controller plasmid regulates expression of *bxb1*, *tp901-1* and *Int5* integrases. The target plasmid corresponds to the 3-input history-dependent scaffold. (**E**) Characterization of a 3-input history-dependent scaffold. We co-transformed the 3-input program with the triple controller plasmid. Bxb1 expression is induced by aTc (input a), Tp901 by arabinose (input b) and Int5 by benzoate (input c). The lineage tree for the program and its corresponding scaffold are represented. For characterizing the system, cells were induced sequentially three times for 16 hours each, with different order-of-occurrences of inputs. Each histogram shows fluorescent reporter expressed in the different states. A representative example is depicted here. Fold change measurements can be found in Figure S12.

We designed five single-lineage 3-input programs and assessed their functionality (Fig. 4E, Fig S11). We performed sequential overnight inductions with aTc (Bxb1), arabinose (Tp901-1) and benzoate (Int5) during 4 days (see methods). The fluorescence measurements by flow cytometry in every input states were consistent with the expected 3-input program (Fig. 4E, Fig. S11). Additionally, the fluorescence fold changes observed for each input state were similar to those observed using OSIRis characterization (Fig. S10, Fig. S12).

We then composed various 3-input programs operating at the multicellular level. We assembled 4 programs (3MP1-4) with different fluorescent reporter genes expressed in response to different input orders in separate lineages (Fig. 5). Bulk fluorescence intensities of all multicellular systems executing 3-input programs were measured by plate reader, and were in good agreement with the corresponding lineage trees (Fig. 5). Here again, the decrease in fluorescence intensity was proportional with the dilution rate of each population in the mix (Fig. S13).

**Figure 5.**
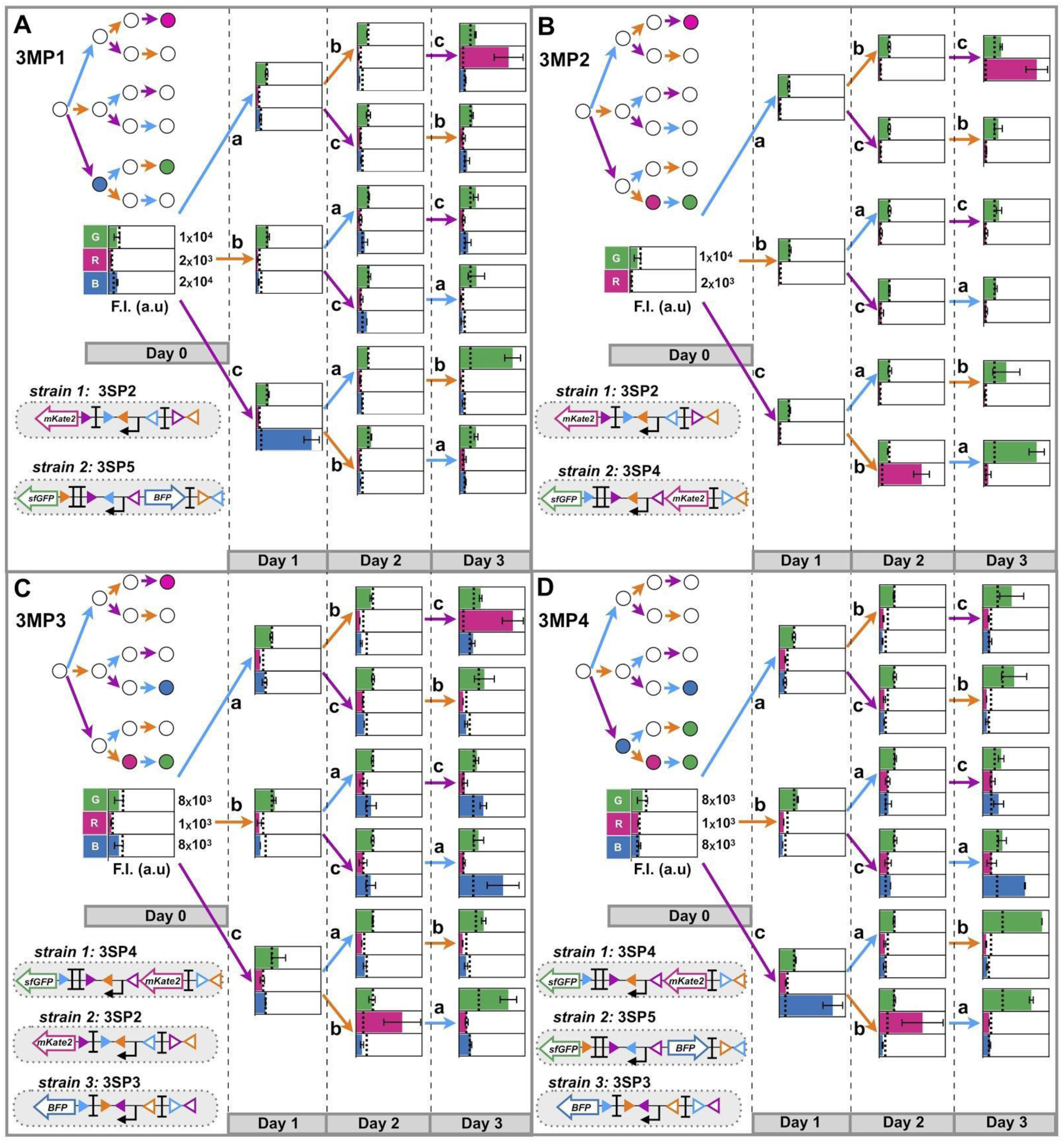
Characterization of 3-input multicellular programs. Design and characterization of 3-input Multicellular Programs operating in various lineages and using different number of strains (**A**-**D**). Programs were decomposed in two sub-programs (**A** and **B**) or three sub-programs (**C** and **D**) implemented in two or three different CCUs, respectively. The inputs are represented by letters, a for aTc (expression of Bxb1 Integrase), b for arabinose (expression of integrase TP901-1) and c for benzoate (expression of Int5). Strains were mixed in equal proportions and grown consecutively during 16 hours, for 4 days in presence of inductor. The bar graph corresponds to the fluorescence intensity (F.I) in arbitrary units (a.u) for each fluorescent chanel (GFP, RFP and BFP), in linear scale. The error bars correspond to the standard deviation for two different experiments measured by plate reader. The dotted line indicates the autofluorescence of negative control strain. Fold change measurements can be found in Figure S13.

Some history-dependent programs also have a combinatorial logic component, i.e. part of the program that can be recapitulated by a Boolean logic device (see sup. Text). We calculated that mixing Boolean and sequential logic devices can actually reduce the number of devices, hence the number of strains used (Fig. S14). We tested this minimization strategy using a 3-input history-dependent program requiring four different strains in fully-sequential mode (Fig. S15A). Combining Boolean and history-dependent devices, only 3 strains instead of 4 are needed to implement the same program (Fig. S15A). The measured fluorescence intensities in mixed populations were consistent with the lineage tree (Fig. S15B), confirming that Boolean-based minimization is indeed a viable approach to reduce multicellular system complexity while obtaining the same cellular program. These results also highlight how the modularity and composability of distributed multicellular computation supports the use of different logic designs within different members of the multicellular system.

We then scaled our scaffold designs and generated scaffolds for 4- and 5-input history-dependent gene expression programs (Fig. S16). The 4-input scaffold allows for expression of a different GOI in each state of a given lineage, while the 5-input scaffold allows expression of a different GOI in each state except in the state 0 (with no input) (Fig. S16B). An additional strain is needed if gene expression is required in this state. Taken together, these data demonstrate the possibility to reliably implement multi-input/multi-output history-dependent programs in a distributed fashion across a cellular population.

### Robustness of history-dependent programs

To evaluate the fidelity of all implemented history-dependent programs, we applied a vector proximity framework (*52*). Using this analysis we calculated the similarity between the biological data for history-dependent programs and their ideal implementation (see methods) (Fig. 6). In this representation, the lower the angle deviation is from 0°, the closer the program behavior is to its expected one. For instance, a program exhibiting a behavior opposite to the expected one would have an angle of 90°. Because the switching rate is essential to obtain a good program implementation, we considered programs with similarity angles smaller than 5° in excellent agreement with the expected outcome; programs with angles between 5 and 10° were qualified as reliable, while the ones with angles higher than 15° were not recommendable to use.

**Figure 6.**
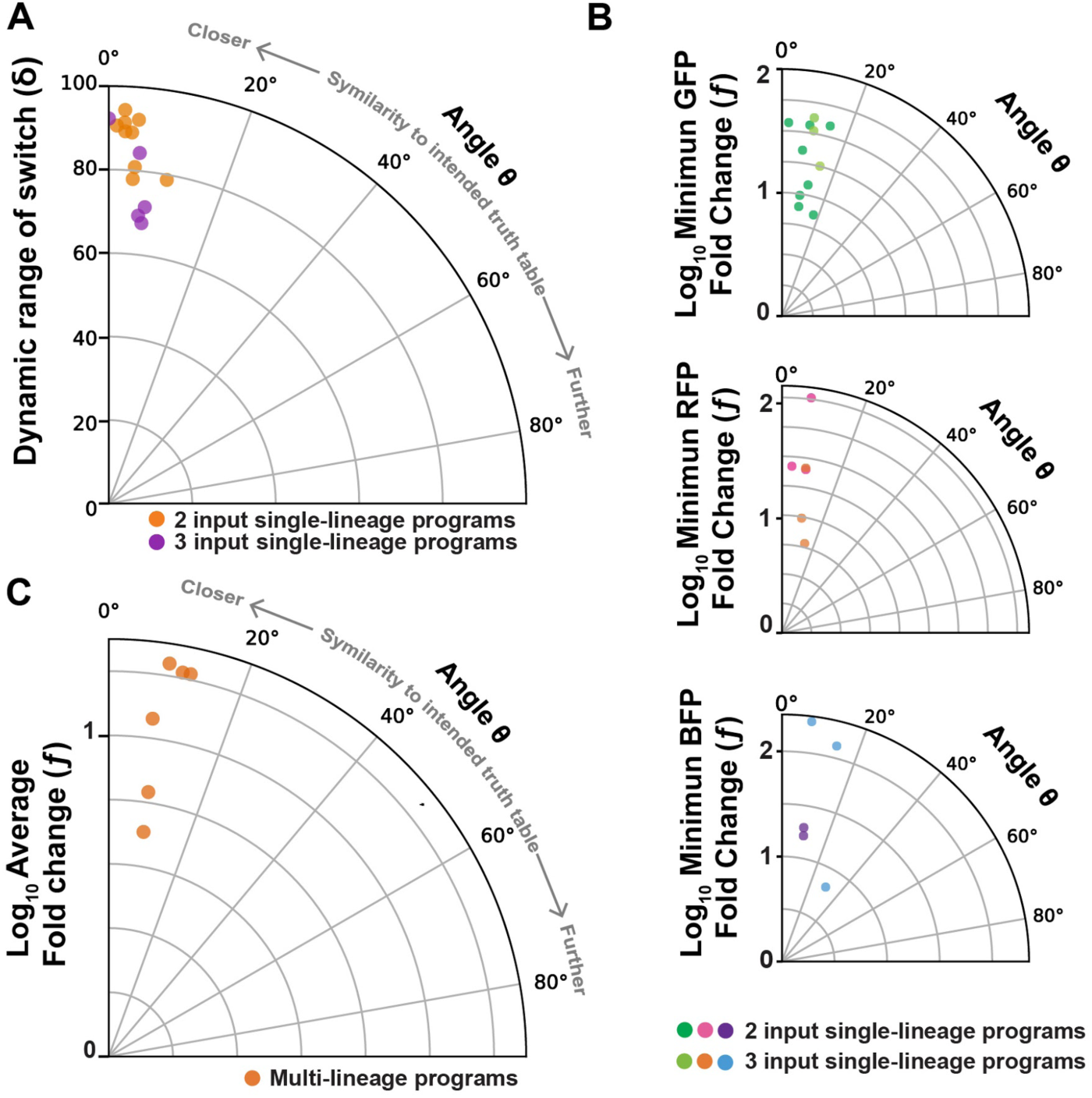
Robustness of history-dependent programs. Angles representing the similarity between biological and the ideal implementation of history-dependent programs were plotted. (**A**) Switching robustness, with angles computed from percentage of cell versus the worst-case switching rate of the switch (□), were plotted. (**B**) Reporter gene expression robustness with angles from fluorescence intensity versus the logarithm of minimum fluorescence fold change (□) for GFP, RFP and BFP, were plotted. The minimum fold change correspond to the fold change between minimum fluorescence in ON input state and the maximum fluorescence in OFF state (□). The *Log_10_* □ were plotted. (**C**) Implemented multicellular programs robustness with angles evaluated between biological and expected fluorescence from plate reader measurement versus the logarithm of average fluorescence fold change were plotted.

We found that single-lineage devices had a very robust switching behavior for both 2- and 3-input programs (Fig. 6A), mostly with angles lower or equal to 5° (11/14) and none resulted in an angle of more than 15°. Because all programs had a high switching rate, we extended our analysis to program output intensities, and monitored the similarity between the experimentally measured fluorescence intensities and the theoretical ones. The robustness of fluorescence was also close to the expected, with angles lower than 17° (Fig. 6B). Only the program 3SP3 presented a higher angle (30°) and lower minimal fold change fluorescence (Table S1), presumably because of a lower BFP expression, as we observed in the fold change analysis (Fig. S12). Additionally, we evaluated the robustness of the implemented multicellular programs by measuring the similarity between expected and experimental fluorescence intensity of the system. We plotted these angles versus the average fold change in fluorescence intensity for each program. We found that the six multicellular programs operated reliably, with angles comprised between 8 to 12 degrees (Fig. 6C). As expected, multicellular programs had fluorescence fold change lower in average than others programs, due to dilution effect. In general, programs behaviors are highly similar to expected ones, demonstrating the predictability of the modular scaffold operation.

### Automated design of history-dependent programs

Automated design frameworks (*19*, *52*, *60*) that lower the entry barrier into novel technologies have proven to be critical in empowering a larger community with technological advances, providing unexpected innovations that could not have been envisioned beforehand. In this context, the automation of genetic circuit design is an important step toward the deployment of cellular computing systems into myriad research or engineering applications.

To automate the design of history-dependent programs, we encoded an algorithm taking a lineage tree as input (equivalent to a sequential truth table) and providing the biological implementation as output (Fig. S17, see methods, and (*61*)). The biological implementation consists of a graphical representation of the genetic circuit and the device DNA sequence of each strain (Fig. 7). To enable broad access to our design framework, we provide a website for systematic and automated design of history-dependent logic called CALIN (Composable Asynchronous Logic using Integrase Networks) (*62*). In the CALIN web-interface, the user fills in the number of inputs to process and the desired lineage tree. The interface provides as an output the DNA architectures of the computational devices and the connection map between inputs and integrases along with the corresponding DNA sequences optimized for *E. coli* which can be directly synthesized to obtain a functional system. The CALIN interface therefore allows the scientific and engineering community to build upon our work to address many different problems.

**Figure 7.**
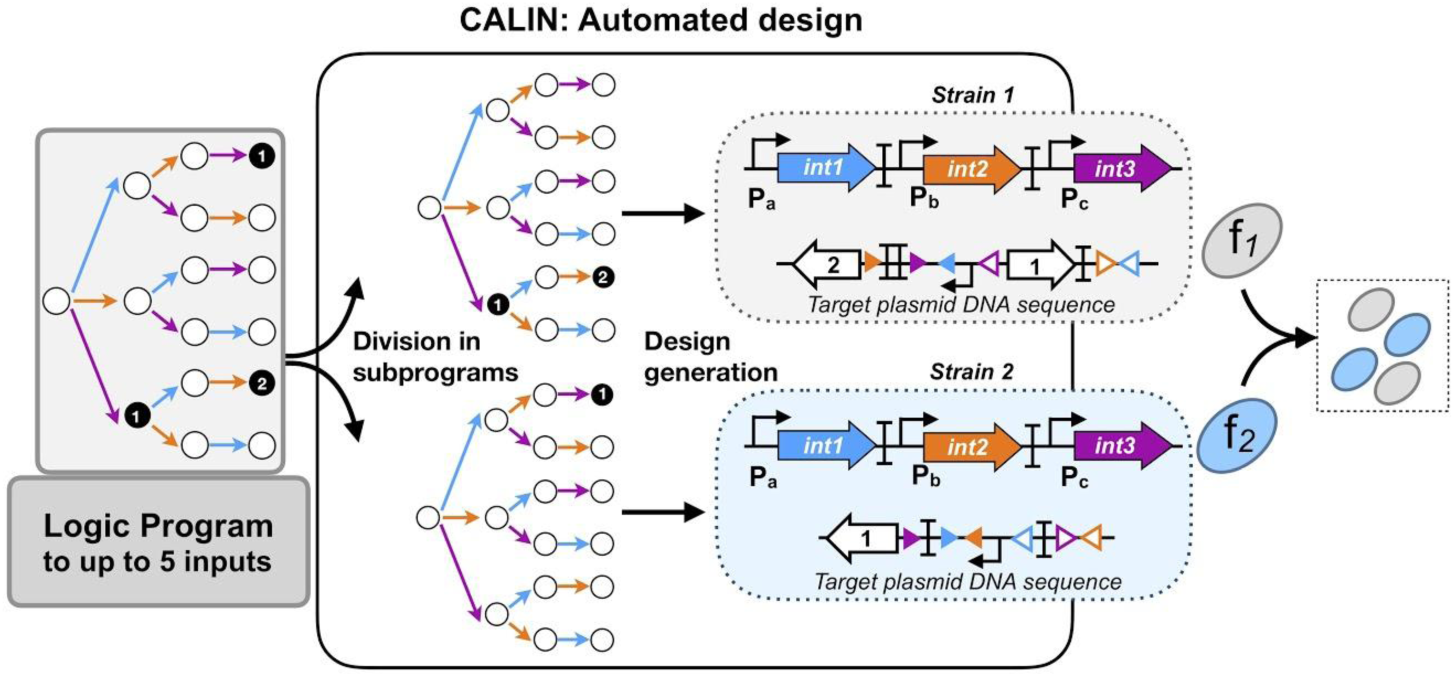
Automated design of history-dependent programs. Automated design of history-dependent programs using the CALIN web interface. The python program takes as input an history-dependent program written as a lineage tree. This program is decomposed in sub-programs, the decomposition is performed by extracting in priority subprograms with ON state at the extremity of the tree (corresponding to state with a high number of inputs). For each subprogram, the algorithm identify the ON states and the order of inputs within the lineage. Based on these informations, the biological design is obtained, with the graphical design of the integrase and the history-dependent devices and its corresponding DNA sequences. By composition of the designs of each subprogram in different strains the full program design is obtained. The CALIN web-interface uses the algorithm, and takes as input the logic program as a lineage tree and gives as output the graphical design and its DNA sequence for each subprogram.

## DISCUSSION

Implementing rich behavior in living organism requires the development of robust frameworks for multicellular history-dependent logic (*63*, *64*). In this work we demonstrate that complex history-dependent programs can be executed in a simple manner by distributed multicellular computation. Multicellular computation provides highly modular and composable systems, *i.e.* many logic functions can be implemented in a straightforward manner from a reduced set of strains executing basic functions. Here we designed, characterized, and optimized 2- and 3-input scaffolds. We built nine 2-input and five 3-input single-lineage logic devices. All devices worked as expected and exhibited predictable behavior close to the ideal program implementation. Without further optimization, we composed various strains containing single-lineage devices to implement complex history-dependent programs executed at the multicellular level. The fact that our system is built by composing well-characterized, standardized genetic elements allowed us to automate program design in a straightforward manner. We provide an easy-to-use web-interface that gives, from a desired program, the corresponding circuit designs and DNA sequences.

The 21 programs tested here represent the largest set of history-dependent programs implemented to date. Using the single-lineage programs from this study, 36 different programs can be executed. We show that 3 states can be unambiguously distinguished using different fluorescent proteins. More states could be distinguished by fluorescence microscopy by fusing fluorescent proteins to subcellular localization tags (*65*) or by expressing various combinations of fluorescent proteins and performing spectral deconvolution (*66*).

Our simple and robust design framework is extendable to four and five inputs. Because intermediate recombination states are encoded within DNA, we developed a rapid prototyping workflow (OSIRiS) based on total synthesis and characterization of all recombination intermediates. We found that the results from OSIRiS characterization closely matched device behavior when operating in real conditions, *i.e.* responding to recombinase mediated inversions or excisions. The OSIRiS workflow is applicable to any recombinase-based circuit.

While we do operate our system in fundamental mode, some inputs could arrive in a mixed manner. The time resolution of the system (i.e. the minimum time between two inputs so that different outcomes can be observed) and the proportion between the resulting populations having entered different lineages would depend on the dynamics of the recombination reactions, which is governed by integrase expression, stability and catalytic rate, among others. In this context, our system could be used to reverse engineer the timing between inputs, as already shown before (*54*).

One potential challenge for application requiring a detectable readout like biosensing is that the system output is produced by a single strain at a time. However, our systems with up to three strains still have a well-detectable output, similarly to previously engineered Boolean recombinase logic systems operating at the multicellular level (*25*). If needed, cell-cell communication could be used to propagate the output state to all the strains of the population (*60*). DNA-sequencing methods could also be used to address the state of the system (*50*, *55*, *67*, *68*). Finally, for many applications like morphogenetic engineering, strain specific phenotypic output would be desirable. By changing output gene identity, various phenotypes like secretion, adhesion, or motility could be implemented in an history-dependent fashion.

Another challenge is the number of strains required to implement programs requiring gene expression in multiple lineages. For example a maximum of 6 strains is required to execute all 3-input programs. Importantly, we showed here and in previous work (Fig. S6 and (*25*)) that multicellular systems operating for three days exhibited reasonable stability. If needed, synthetic cooperative behavior could be used to maintain all strains in the population (*9*, *10*). Strains could also be cultivated in separate chambers within a microfluidic device for instance, avoiding competition problems. Alternatively, the number of strains required to implement complex logic circuits can be reduced by combining Boolean (*25*) and history-dependent devices. Future research exploring minimization strategies and alternative scaffold designs could help reduce even more the number of required strains.

The functionalities of our system could be expanded in several manners. For instance, while integrases mediate irreversible switching, the addition of recombination directionality factor (RDF) (*49*) would result in history-dependent programs with reversible state transitions and increased complexity. Cell-cell communication could also provide another layer of complexity to coordinate synthetic collective behavior (*69*).

Because of the high-modularity of integrase logic, the system presented here could be quickly repurposed by swapping the signals controlling integrase expression to detect chemical (e.g. biomarkers of disease, pollutants) or physical signals (e.g. temperature, light). Here we show using the benzoic acid sensor BenR that a viable approach to engineer new switches is to start by optimizing the properties of the transcriptional controller and then screen for different integrase variants with different translational efficiency or stability (Fig. S18). As serine integrases work in many species, including mammals and plants, the history-dependent programs presented here could be used in myriad research and engineering applications. Programs responding to environmental signals could enable therapeutic cells to act in a highly precise spatiotemporal fashion, increasing specificity and therapeutic efficacy while reducing side effects. Cellular sensors could perform temporal logic analysis for diagnostics, environmental, or manufacturing applications. Finally, sequential programs would support iterative, ordered assembly of synthetic biological architectures for tissue and biomaterial engineering (*70*, *71*).

## Acknowledgments

We thank the synthetic biology group and members of the CBS for fruitful discussions.

## Author contributions

S.G., A.Z., and J.B designed the project. S.G., A.Z., Z.B.M., M.C., and P.M. performed the experiments. S.G., A.Z., Z.B.M., M.C., L.C., and P.H. analyzed data. S.G. wrote the python source code for CALIN and V.M., S.G. implemented the web server for CALIN. S.G., A.Z., L.C., P.H. and J.B. wrote the manuscript.

## Data and materials availability

DNA sequences for all constructs are available as a supplementary file.. All raw data are available in supplementary materials. All codes, and materials used in the analysis are available in supplementary materials and on github. Plasmids are available from addgene: 2SP1 ID:126526; 2SP2 ID: 126527; 2SP3: 126528; 2SP4 ID 126529; 2SP5 ID: 126530; 2SP6 ID: 126531; 2SP7 ID: 126532; 2SP8 ID: 126533; 2SP9 ID: 126534; 3SP1 ID: 126535; 3SP2 ID: 126536; 3SP3 ID: 126537; 3SP4 ID: 126538; 3SP5 ID: 126539; pITC (Integrase triple controller) ID: 126540.

## Materials and methods

### Strains, media, and inducers

*E. coli* strain DH5alphaZ1 (laci^q^, PN25-tetR, Sp^R^, deoR, supE44, Delta(lacZYA-argFV169), Phi80 lacZDeltaM15, hsdR17(rK-mK+), recA1, endA1, gyrA96, thi-1, relA1) was used in experimental measurements of history-dependent programs. DH5alphaZ1 was grown on LB media with antibiotic corresponding to the transformed plasmid(s) to do the cloning. For experimental measurements the cells were grown in Azure Hi-Def medium (Teknova, 3H5000) supplemented with 0.4% of glycerol. Antibiotics used were kanamycin 25μg/mL and chloramphenicol 25μg/mL. The inducers were: L-arabinose (Sigma-Aldrich, A3256) used at final concentration of 0.7% wt/vol; anhydrotetracycline (Sigma-Aldrich, 37919) used at final concentration of 20 ng/mL and sodium benzoate (Sigma-Aldrich, B3420) used at final concentration of 100 μM, for 3-input programs.

### Integrase controller and target plasmids construction

One-step isothermal Gibson assembly was used (*72*) to build all plasmids described. Vectors pSB4K5 and J64100 (from parts.igem.org) were used to construct all genetic circuits. The pSB4K5 vector containing kanR cassette and pSC101 origin of replication was used to clone different input programs, based in DNA scaffold design, including BP and LR targets, genes of interest and others DNA parts. The J64100 plasmid containing camRY cassette and ColE1 origin of replication was used to clone the integrase controller cassette. Enzymes for the one-step isothermal assembly were purchased from New England BioLabs (NEB, Ipswich, MA, USA). PCR were performed using Q5 PCR master mix and One-Taq quick load master mix for colony PCR (NEB), primers were purchased from IDT (Louvain, Belgium), and DNA fragment from Twist Bioscience. Plasmid extraction and DNA purification were performed using kits from Biosentec (Toulouse, France). Sequencing was realized by GATC Biotech (Cologne, Germany).

To build the triple controller plasmid, we constructed an inducible cassette containing the *benR* gene regulator, constitutively expressed by promoter J23106, the P_ben_ promoter (both *benR* and P_ben_ from metabolic and sensing modules, respectively (*73*)) and Integrase 5 (*57*) controlled by P_ben_ promoter (see vector map). The cassette was inserted upstream of duall controller integrase plasmid (*49*). After assembly the triple controller vector was transformed and cloned in *E. coli* strain DH5alphaZ1.

The design of DNA scaffold for target plasmid was done using a python script to automatically generate a library of DNA sequences (https://github.com/synthetic-biology-group-cbs-montpellier/Generate_DNAseq), which minimizes the number of errors. Moreover, because the final sequences result from permutations of a reduced set of parts, Python is particularly well suited for the task. All sequences were designed to support cloning by Gibson assembly at an identical location in pSB4K5 template vector. Consequently, all sequences were composed of the 40 bp spacer 0 at 5’ end, and 40 bp spacer N at 3’ end. The DNA sequences for every designed program were synthesized, as linear fragments, by Twist Bioescience. Each DNA fragment was PCR amplified and assembled between spacer 0 and N in pSB4K5 template vector. All DNA sequences of history-dependent programs are listed in Table S3. Target plasmids were transformed and cloned in *E. coli* strain DH5alphaZ1. All plasmids were purified using QIAprep spin Miniprep kit (Qiagen) and sequence verified by Sanger sequencing in Eurofins Genomics, EU.

### Experimental conditions and sequential induction assays

Integrase controller and target plasmids were co-transformed in *E. coli* strain DH5alphaZ1 and plated on LB agar medium containing kanamycin and chloramphenicol antibiotics. Three different colonies for each program to test were picked and inoculated, separately, into 500μL of Azure Hi-Def medium (Teknova, 3H5000) supplemented with 0.4% of glycerol, kanamycin and chloramphenicol in a 96 DeepWell polystyrene plates (Thermo Fisher Scientific, 278606) sealed with AeraSeal film (Sigma-Aldrich, A9224-50EA) and incubated at 30°C for 16h with shaking (300 rpm) and 80% of humidity in a Kuhner LT-X (Lab-Therm) incubator shaker. All experiments were performed in the same condition of growth. After overnight growth the cells were diluted 1000 times into fresh medium with antibiotics and let them growth at 37°C for 16h (Day 0). For multicellular programs, cell strains harboring different programs, were equally mixed, before to dilute (1000-fold total dilution) into fresh medium with antibiotics. The mixed populations were grown at 37°C for 16h (Day 0). After 16h of incubation the cells were serial diluted (1000-fold total dilution), first 10μL culture into 190 μL of fresh medium in presence of antibiotics and a second dilution (from first dilution) 10 μL into fresh media with antibiotics and inducers (aTc, Arabinose or Benzoate) or not. The cells were grown at 37°C for 16h (Day 1). After overnight incubation the cells were serial diluted (1000-fold total dilution), first 10μL culture into 190 μL of fresh medium in presence of antibiotics and a second dilution (from first dilution) 10 μL into fresh media with antibiotics and inducers (aTc, Arabinose or Benzoate) or not. (Day 2, aTc→Ara; Ara→aTc, for 2-input programs and aTc→Ara; aTc→Benzoate; Ara→aTc; Ara→Benzoate; Benzoate→aTc; Benzoate→Ara, for 3-input programs). For 3-input programs, after overnight incubation the cells were serial diluted (1000-fold total dilution), first 10μL culture into 190 μL of fresh medium in presence of antibiotics and a second dilution (from first dilution) 10 μL into fresh media with antibiotics and inducers (aTc, Arabinose or Benzoate) or not. (Day 3, aTc→Ara→Benzoate; aTc→Benzoate→Ara; Ara→aTc→Benzoate; Ara→Benzoate→aTc; Benzoate→aTc→Ara; Benzoate→Ara→aTc). To measure cell fluorescence, aliquot of cells from each day were diluted 200 times into Attune Focusing Fluid (Thermo Fisher Scientific, A-24904) and incubated for 1 hour at room temperature before flow cytometry analysis. For plate reader measurement an aliquot of cells from each day were diluted 4 times in phosphate buffered saline PBS before read. For microscopy analysis aliquots of cells from each day were mixed with glycerol at final concentration 15% v/v and kept at 80°C until its analysis. Every experiment was repeated 2 or 3 times.

### OSIRiS design and characterization

To automatized the design of intermediate recombination state, we developed a python script giving from any integrase-based scaffold and a list of integrase of interest the recombination intermediate DNA sequences. OSIRiS code is widely available on github (*58*), requires Python 2.7 and biopython installation.

To use OSIRiS script, a csv file containing integrase sites of interest is required, we provided a csv file containing widely used integrase sites but any specific sequences can be added. Of note, this script is only considering irreversible DNA excision and inversion. As output, genbank file with integrase site annotations is obtained for all intermediate recombination states.

In our scaffold design, we selected the fluorescent proteins sfGFP, mKate2, and BFP, as their excitation and emission spectrums do not overlap. We used P6 as the promoter and B0034 as the ribosome binding site. To insulate the translation from the genetic context, we placed a ribozyme in 5’ end of each output gene, catalyzing the cleavage of the mRNA at this position (*74*). We used different ribozymes for each output gene (RiboJ, BydvJ, and AraJ) to avoid multiple repetitions of sequences in the construct. Based on this design, we generated intermediate recombination input states using our OSIRiS python script. We then synthesized and constructed these sequences. We characterized all the constructs by flow cytometry. The fold change over the negative control was determined from mean value over that of the negative control. The mean fold change was represented in the figure corresponding to the mean of the fold change of the three different experiments.

### Flow cytometry and plate reader analysis

Flow cytometry was performed on Attune NxT flow cytometer (Thermo Fisher) equipped with an autosampler and Attune NxT™ Version 2.7 Software and BD LSR Fortessa (Becton Dickinson), with FACSDiVa software. Experiment on Attune NxT were performed in 96 well plates with setting; FSC: 200V, SSC: 380V, green intensity BL1: 460V (488 nm laser and a 510/10 nm filter), red intensity YL2: 460V (561 nm laser and a 615/25 nm filter). Setting for experiments on Fortessa were FSC: 400V, SSC: 300V, green intensity GFP: 580V (488 nm laser and a 530/30 nm filter), red intensity mCherry: 565V (600nm laser and a 610/20 nm filter) and blue intensity V1: 460V (405nm laser and a 450/50 nm filter). All events were collected with a cutoff of 20,000 events. Every experiment included a positive control expressing GFP, RFP or BFP and a negative control harboring the plasmid but without reporter gene, to generated the gates. The cells were gated based on on forward and side scatter graphs and events on Single cell gates were selected and analyzed, to remove debris from the analysis, by Flow-Jo (Treestar, Inc) software.

Plate reader measurements were done on Cytation 3 microplate reader (Biotek Instruments, Inc). After induction time cultures were diluted four times in PBS and measured with the following parameters (GFP: excitation 485 nm, emission 528 nm, gain 80, BFP: excitation 402 nm, emission 457 nm, gain 70, RFP: excitation 555 nm, emission 584 nm, gain 100, absorbance: 600 nm). For each sample, GFP, BFP, and RFP fluorescence intensity were normalized to absorbance at 600 nm. The arbitrary units of fluorescence were used to graph bars in each figure. Plots for all data analysis were obtained using phyton plotting codes available on github (*75*)

### Microscopy and microfluidic analysis

Cells samples from sequential induction experiment were analyzed by confocal microscopy. Images were acquired using an Zeiss Axioimager Z2 apotome, Andor’s Zyla 4.2 sCMOS camera (MRI platform, montpellier). 2 μL of cells were spotted on Azure/glycerol 2% agarose pads. Images were taken from phase contrast, GFP, RFP and BFP fluorescence images at a 100X magnification. Images were analyzed using OMERO software. Source images are provided as a *Source Data file*.

For microfluidic analysis experiments were performed using a mother machine microfluidic device consisting in arrays of parallel chambers (1µm x 1µm x 25µm) connected to a large channel. Chambers were fabricated using electron-beam lithography on SU-8 photoresist (MicroChem) while the channel was fabricated using soft-lithography. From the subsequent master wafer, microfluidic chips were molded in polydimethylsiloxane (PDMS) and bonded to a glass slide using plasma activation. Cells, grown overnight in LB supplemented with Cam and Kan, were then loaded into the chambers by centrifugation on a spin-coater using a dedicated 3D printed device. LB media flown in the mother machine is supplemented with Cam and Kan, but also with 5 g l^−1^ F-127 pluronic to passivate the PDMS surfaces and prevent cell adhesion. The medium diffuses to the chambers, providing nutrients and chemical of interest to cells. Chemical inducers (aTc at 200ng/mL and Arabinose at 1%) were added to the media as required using solenoid valves (The Lee Company). A peristaltic pump was used to flow the various medium through the device at a flow rate of 90 µl/min. Both the microfluidic device and the medium were constantly held at 37 °C. Images were obtained using an inverted Olympus IX83 microscope with a 60x objective. Fluorescence levels were measured within a small rectangular region of interest located at the top of each chamber where a single cell is trapped.

### Automated generation of genetic designs

#### Equations for the determination of number of subprograms to implement in different strains for history-dependent logic

History-dependent programs are represented as a lineage tree. Each node of this tree corresponds to a specific state of the system in response to a different scenario: when no input occurred, when one input occurred, and when multiple inputs occurred in a particular sequence. For an N-input program, the number σ of states is equal to

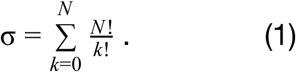

Then, for N-input/1-output history-dependent logic programs, the number *P* _1_ of possible programs is equal to 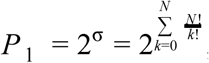, as all states can have either a ON or OFF output. Similarly for N-input/M-output history-dependent logic programs, 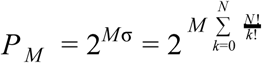 programs exist.

The maximum number of strains needed to implement an N-input/M-output history dependent gene expression program is equal to N!, which corresponds to the number of possible lineages in an N-input lineage tree.

#### Automated generation of genetic designs to execute multicellular history-dependent gene expression programs

We encoded an algorithm capable of creating up to 5-input history-dependent program design using Python (Fig. S17) (*61*, *62*). The algorithm takes as input a lineage tree (equivalent to a sequential truth table). The output corresponds to the biological implementation, such as a graphical representation of the genetic circuit and its associated DNA sequences for each strain. The lineage tree is decomposed into sub-trees consisting of a single lineage containing one or multiple ON states. This decomposition is done by iteratively subtracting the lineages containing ON states. To obtain the lowest number of sub-programs, we prioritize among the lineages with ON states the ones for which the highest number of inputs occurred (from the right to the left of the lineage tree). After decomposition, for each selected lineage, two pieces of information are extracted. First, based on which states are ON, we directly design the corresponding scaffold by specifically inserting genes at the adequate GOI positions. Second, the order-of-occurrence of inputs corresponding to the lineage is used to identify which sensor modules are needed among the different connection possibilities between control signals and integrases. Then, by combining the design of the different lineages, we obtain the global design for biological implementation of the desired history-dependent gene-expression program. To simplify the construction process of logic circuits, DNA sequence of computation devices is generated by our Python code. In CALIN, sequences are adapted for *E.coli*. But sequence generation can be adapted to other organisms (databases are available for *Bacillus subtilis* and *Saccharomyces cerevisiae*) or customly designed using the source Python code available on github (*61*)

### Robustness analysis

We adapted and generalised the vector proximity method described in Weinberg et al. 2017 (*52*) in order to compare the outputs of the programs to the desired theoretical behaviour.

Each implemented program has a different number of inputs (up to 3) and outputs; the matrix *T* corresponding to the expected outcomes presents a number of rows equal to the number of states σ, Eq.(1), and three columns maximum (the RGB output channels). This matrix can be mapped to a 3σ-dimensional vector *t* by stacking all rows of *T* one on top of the others. Similarly, we constructed a vector *s* from the table *S* of the experimental outcome (percentage of switched cells or fluorescence values for each channel and state). From the definitions of the scalar product 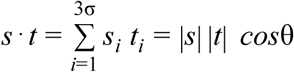, we can compute the angle θ between *s* and *t* as 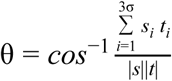, where *s_i_* and *t_i_* are the i-th components of the vectors *s* and *t*, and |·| represents the Euclidean norm. If *s* and *t* are proportional, i.e. if the experimental output is an ideal implementation of the program, the angle is 0°. The larger the angle, the worse the accuracy of the program with respect to the desired behaviour.

As explained below, the table *T* gathering the expected program’s outcomes is computed differently when comparing percentage of switch or fluorescence from a single CCU, or when testing a program composed of many CCUs.

When analysing the percentage of switch for each state from flow-cytometry data (Fig. 6A), the values in the signal table *S* are bound by 1 (when all cells switch), and the table *T* is the actual binary truth table. Instead, when comparing fluorescence values for single strains (1 CCU) as in Fig. 6B, the 1’s of the table *T* are replaced by the fluorescence of the positive control (measured by flow cytometry) of each corresponding reporter gene (referred to strains expressing constitutively the fluorescence reporter gene), and the 0’s by the negative control (background values). This allowed us to consider fluorescence variation in programs with different GFP variants, and to avoid capping the signal values as done in Weinberg et al. 2017. In case of multicellular systems (several CCUs), the table *T* is constructed by summing the fluorescence measured with the plate reader for each individual CCU, then the resulting values are divided by the total number of CCUs used to implement the multicellular program (Fig. 6C). This procedures relies on the hypothesis that each strain equally contributes to the total observed fluorescence. The matrix *T* gathering the expected outcomes can then be compared with the experimental output *S* as previously explained.

We quantified the strength of the signal output by computing the dynamic range δ of switch, considering the absolute difference between the minimum percentage of cells in the ON input-state and the maximum percentage of cells in the OFF input-state (radial coordinate in Fig. 6A). In mathematical terms, we computed

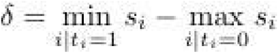

When analysing fluorescence values of individual CCUs (Fig. 6B), we calculated the minimum fold change *f* of each program considering the fold change between minimum fluorescence in the ON input-state and maximum fluorescence in the OFF input-state for each RGB channel:

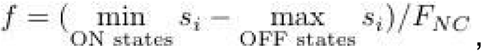

bearing in mind that the matrix *S* in this case gathers fluorescence values and where *^F^_NC_* is the fluorescence of the negative control for the corresponding output channel. In the radial coordinate of Fig. 6B we plotted the *Log_10_ f*, meaning that a radial coordinate of 1 corresponds to a fold change *f=10*.

Finally, we average the fold change of different channels when multiple CCUs are used in the implementation of multicellular programs and plot the logarithm of its value (Fig. 6C).

## Supplementary Text

**Supplementary Text S1. Minimization of history-dependent circuits using Boolean logic devices**

The number of strains required for implementing history-dependent gene-expression programs can be reduced using Boolean logic devices. Indeed, gene-expression programs independent of the history of occurrence of inputs are implementable using Boolean logic (*60*). Some history-dependent gene-expression programs are decomposable into Boolean logic function(s) and history-dependent subprogram(s). The combination of history-dependent and Boolean logic devices allows a reduction in the number of strains required for the implementation of some history-dependent gene-expression programs. We created an algorithm in Python to automate this simplification. We generated all Boolean functions and converted each truth table into a lineage tree. To implement history-dependent programs, we tested if any Boolean functions can be extracted from this program. If the use of Boolean devices leads to an implementation with an equal or reduced number of strains, the design is saved. We then obtained as output a list of designs based on Boolean and/or history-dependent devices implementing the input program with the minimal number of strains possible. We applied this brute-force method to all 3-input/1-output programs, totaling 65,536 programs. This strategy allows for a reduction in the number of strains for the implementation of 20% of these programs. The results did not significantly reduced the median number of strains required for the implementation of history-dependent programs, however 48% of 3-input/1-output programs are decomposable using Boolean programs while minimizing the number of strains (Fig. S14).

**Figure S1.**
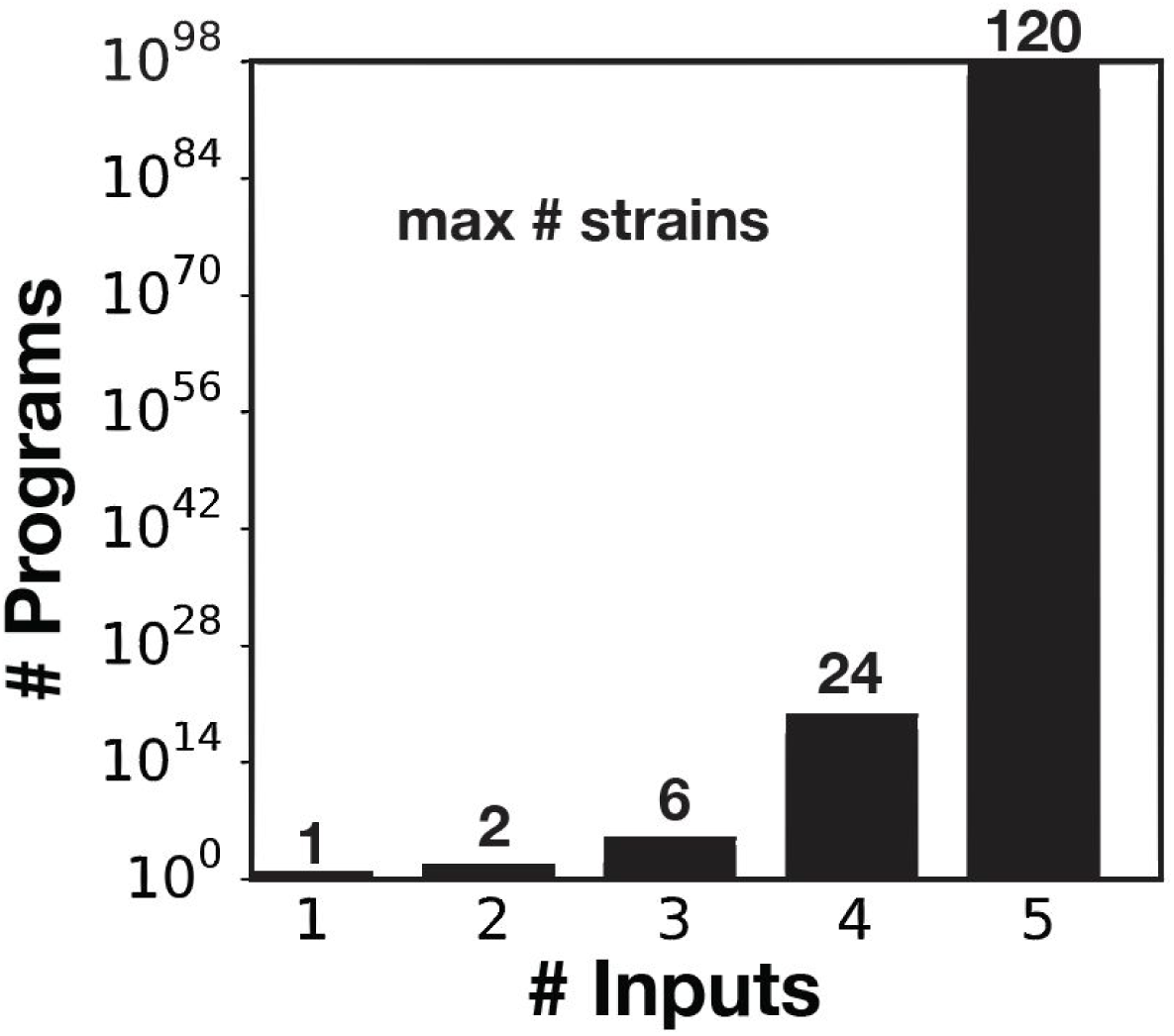
Number of cellular computation units needed to implement a history-dependent programs. Number of single-output history-dependent programs and maximum number of strains needed for 1 to 5 inputs. The bar graph represents the number of single-output history-dependent programs from 1 to 5 inputs and the number at the top of each bar corresponds to the maximum number of strains required for the implementation of these programs. See materials and methods for detailed equations.

**Figure S2.**
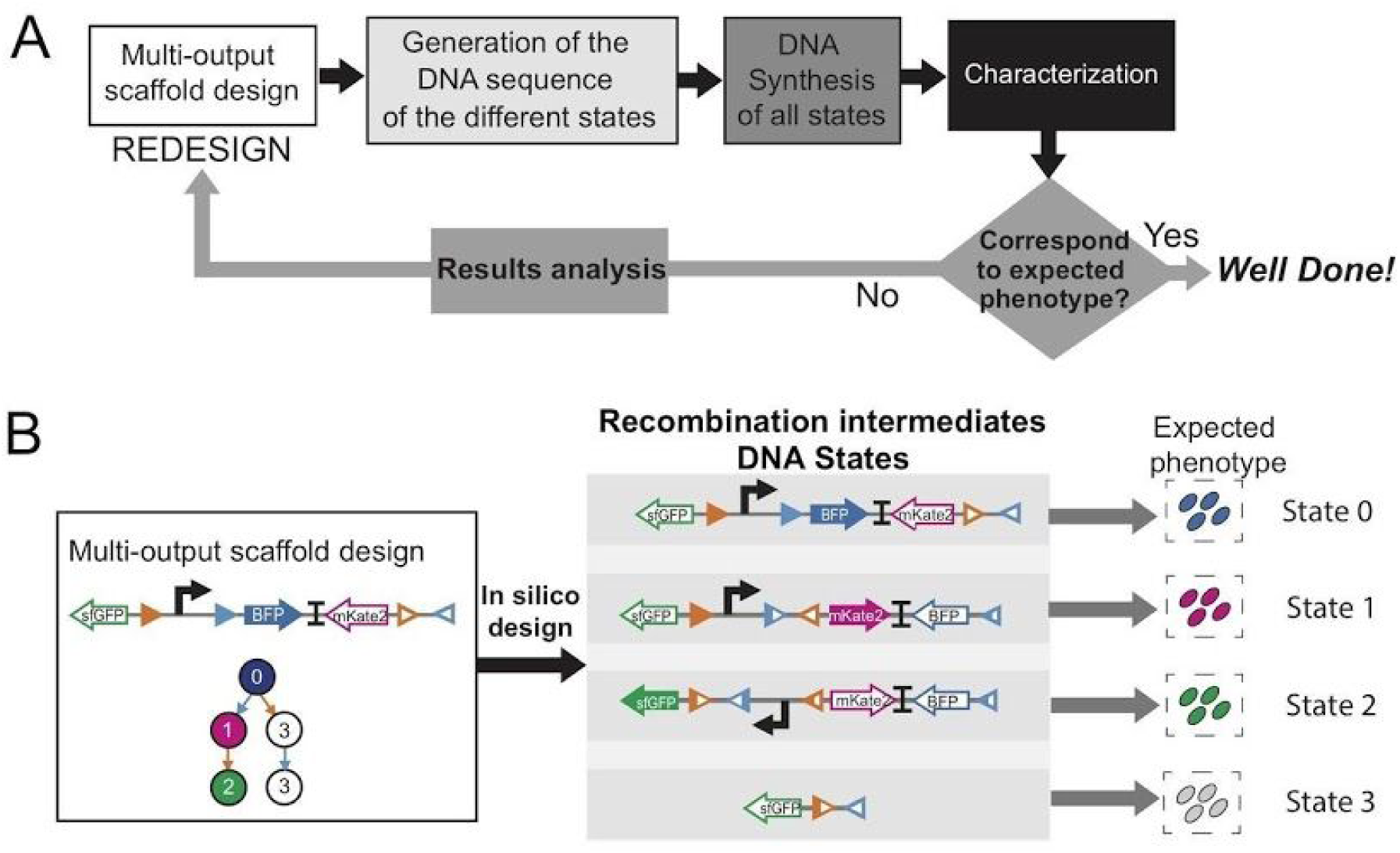
Optimization by Synthesis of Intermediate Recombination States. **(A)** OSIRiS workflow. A history-dependent scaffold corresponding to a specific lineage and with expression of a different gene in each input state is designed. The DNA sequences of the different input states corresponding to the intermediate recombination states are generated (DNA states). The initial and the intermediate sequences are synthesized and characterized and the phenotypes are compared to the expected phenotypes. If they match, a celebration is performed. Otherwise the results are precisely analyzed to identify the origin of the bug, the multi-output scaffold is redesigned and a new OSIRiS cycle is performed. **(B)** 2-input OSIRiS workflow. For 2-input program, a scaffold with consecutive expression of BFP, RFP and GFP in one lineage is designed. From this design, the 3 intermediate DNA states are generated and the expected phenotype for each of the 4 DNA states of the entire lineage tree is predicted.

**Figure S3.**
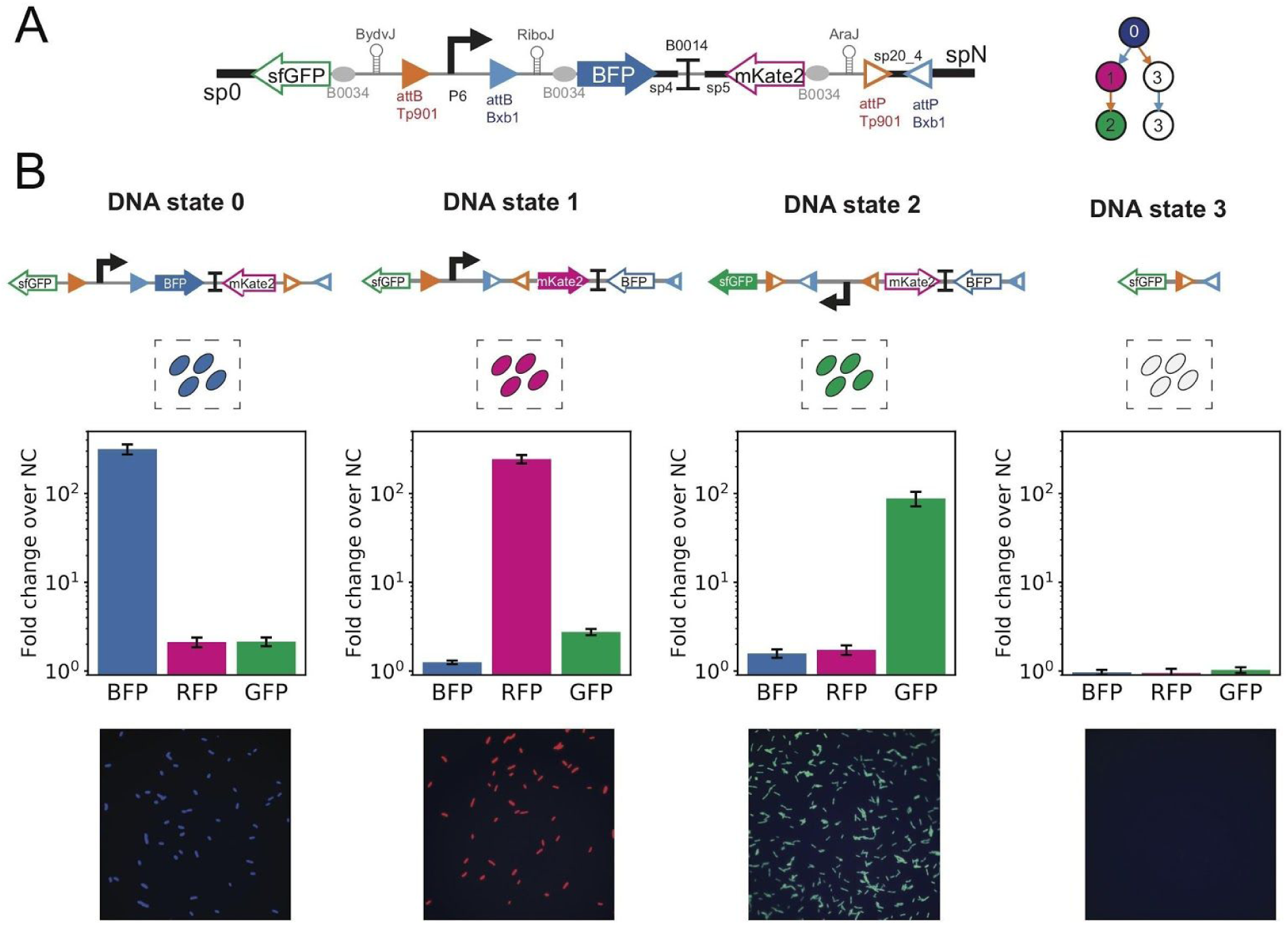
Design and characterization of 2-input OSIRiS. **(A)** Detailed design of the 2-input scaffold for the lineage A then B with Bxb1 for input A and Tp901 for input B. As output gene, we used BFP, RFP and GFP and in 5’UTR of each gene we placed a ribozyme and B0034 RBS to have a translation isolated from genetic context. We used P6 as promoter and B0014 as bidirectional terminator between BFP and RFP coding sequences. We added 40bp spacers for gibson assembly (sp0, sp4, sp5 and spN) and a 20bp spacer between to juxtaposed integrase sites (sp20_4). **(B)** Characterization of the 2-input OSIRiS by flow-cytometry. We characterized each initial and intermediate recombination states in a flow-cytometry by measurement of GFP, RFP and BFP fluorescence intensities. The bar graph corresponds to the fold change over the negative control (strain without fluorescent protein) for each channel. The error bar corresponds to the standard deviation between the fold change obtained from 2 biological replicates. The microscopy images correspond to merge images of the GFP, RFP and BFP channels.

**Figure S4.**
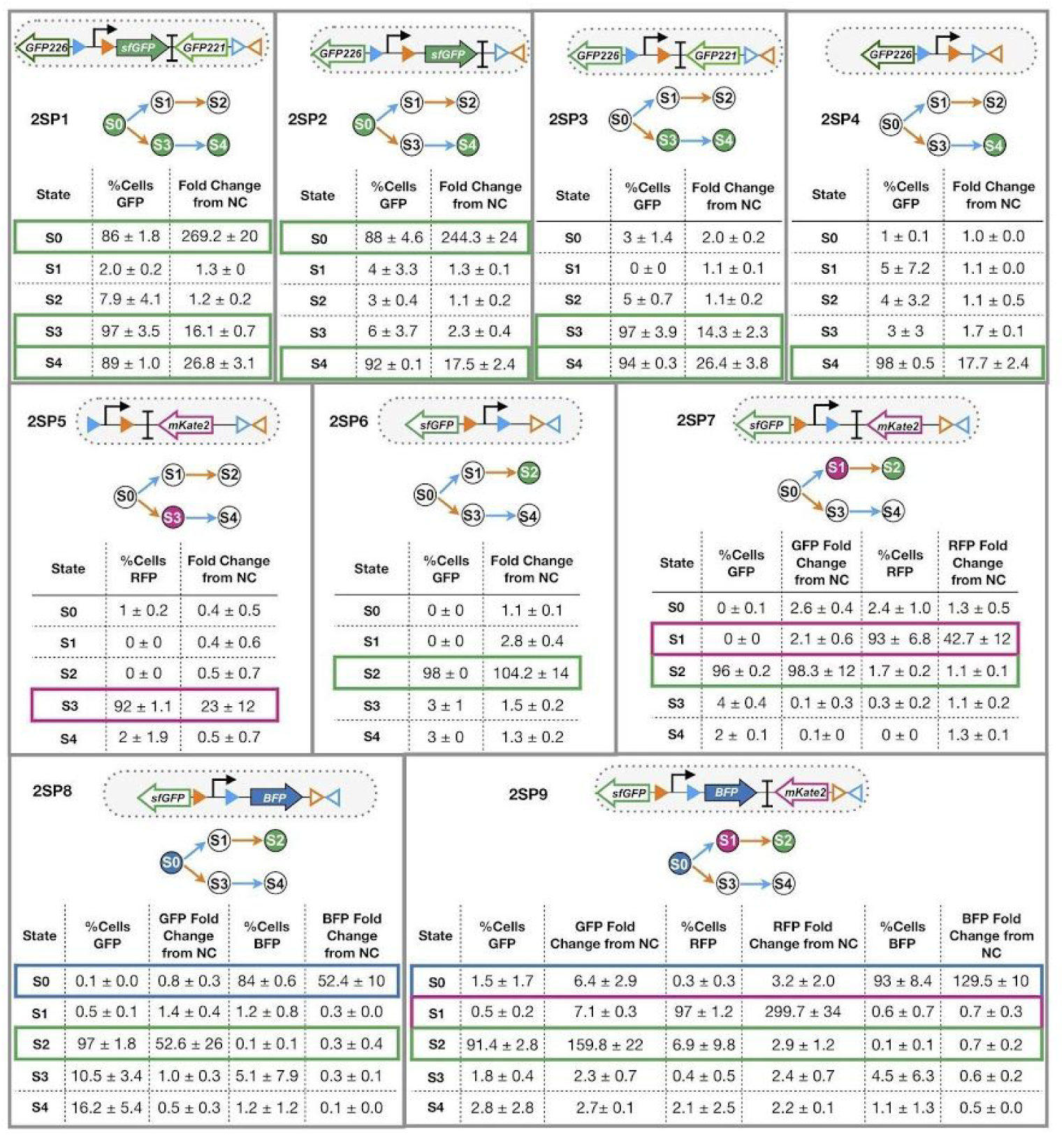
Percentage of cells and fluorescence fold change in each different input states for 2-input single-cell programs. The lineage tree for each program and its corresponding genetic diagram are represented. Each table shows % of cell and reporter fluorescence in gates for respective fluorescence channel, as a result of different induction condition corresponding to an input state of the lineage tree. Values correspond to averages and standard deviations between two different experiments, with tree different colony-forming units analyzed for experiment.

**Figure S5.**
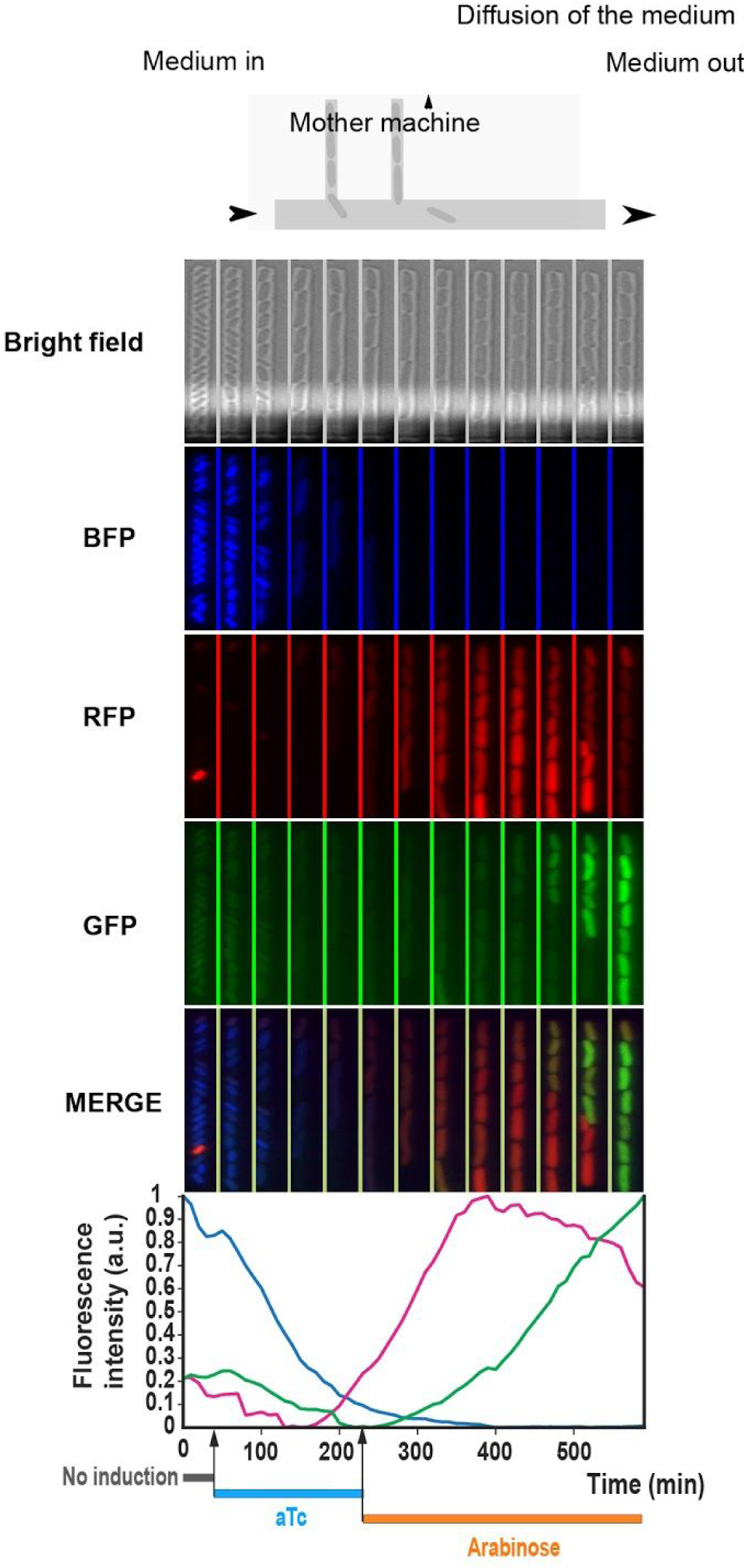
Time needed between inputs in 2-input programs. Cells were grown in a mother machine microfluidic chamber for 10 h. They were grown for 30 min without any induction. After that the induction with aTc was performed for 3 h. Next, the medium was changed and then the Arabinose induction was performed. Microscopy images show the cell fluorescence expression in the mother machine during this time. The graph represents the kinetics of fluorescence intensity normalized over the maximum fluorescence intensity for each channel, thus for each reporter gene in cells exposed to different inducers in order corresponding to the scaffold lineage.

**Figure S6.**
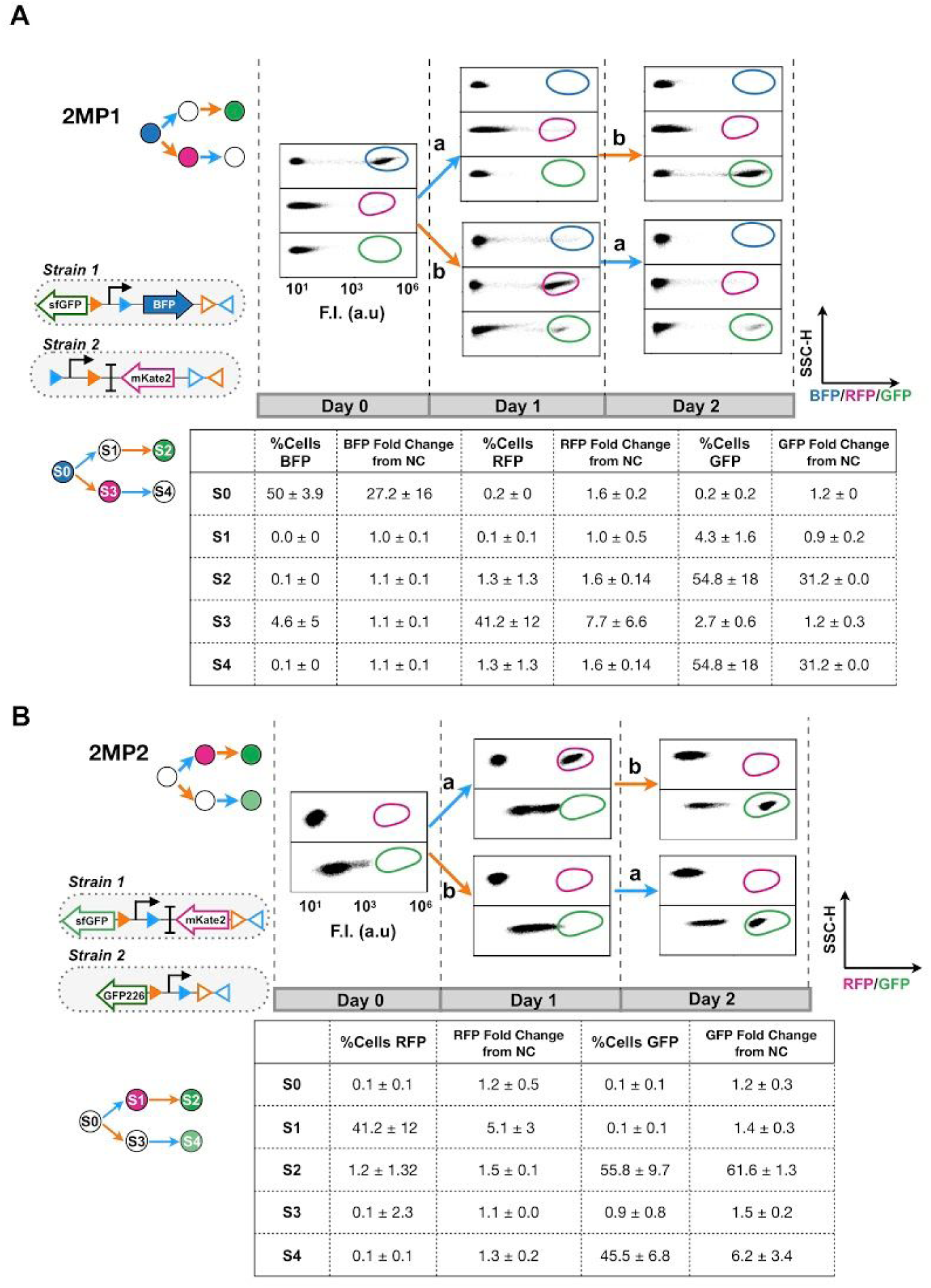
Dot plot from lineages of 2-input multicellular programs. Flow cytometer analysis of populations in 2-input multicellular programs 2MP1 **(A)** and 2MP2 **(B)**, with statistical data of fluorescence fold change and cell percentage on fluorescence channel, for each subpopulation in different input states. The corresponding standard deviation is shown. Scatter plots showing SSC-H (side scatter height) versus fluorescence intensity (F.I), are representative of two independent experiments, with three different replicates. Channels for BFP, RFP and GFP, from top to bottom are plotted. Positive controls expressing fluorescence reporter genes were used to set the gates for each fluorescence channel.

**Figure S7.**
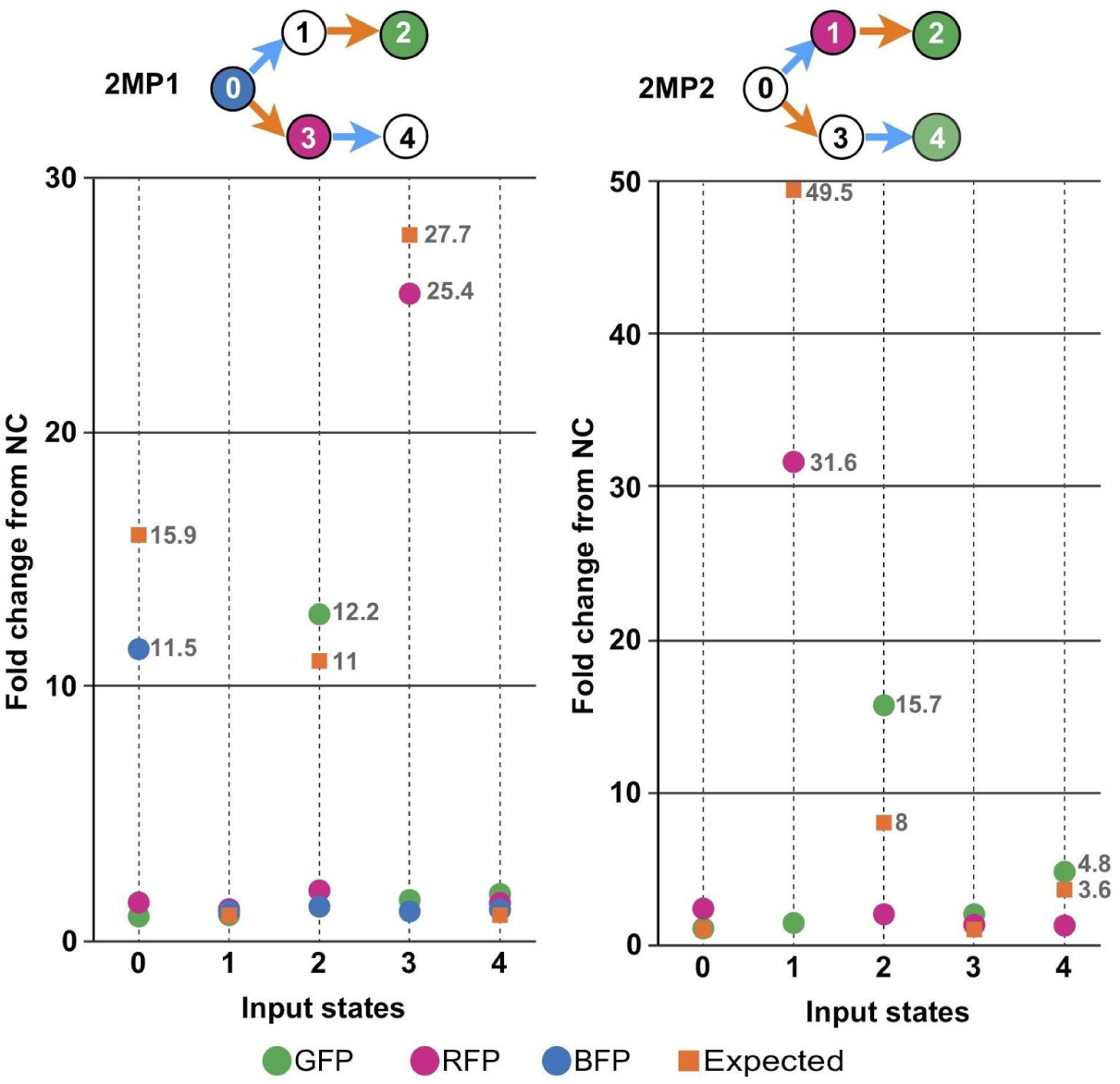
Fold change differences between resulted and expected values for 2-input multicellular programs. Plots show the fold change fluorescence over the negative control, resulted for 2-input multicellular programs in every input states. The expected value was obtained dividing single cells fold change fluorescence values by the number of strains which compose the multicellular system.

**Figure 8. Figure S9.**
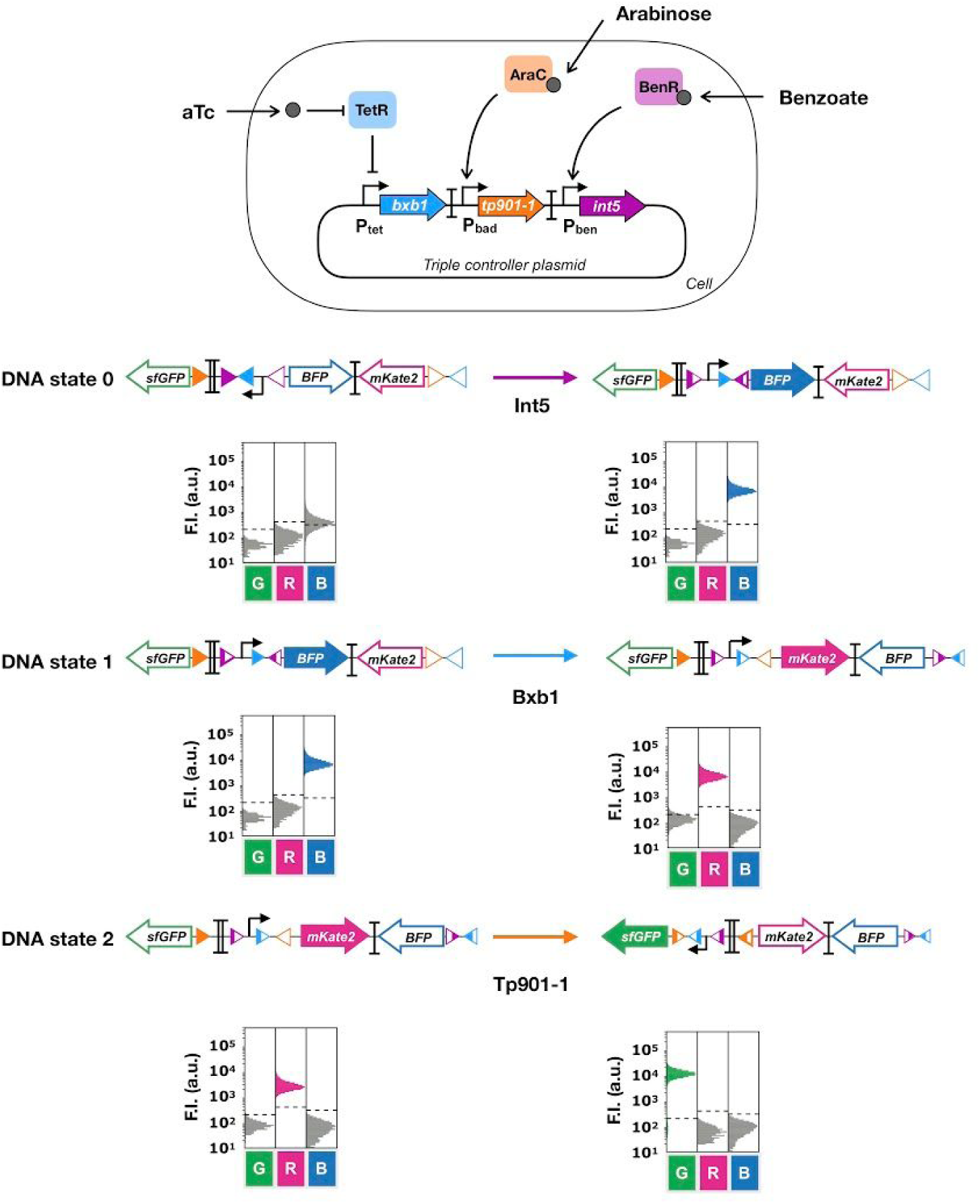
Test of recombination activity of triple controller. Cells harboring the triple controller plasmid were transformed with OSIRiS states for 3-input scaffold, to test the recombination activity of Integrase 5, Bxb1 and Tp901-1 integrases. Each line corresponds to the transition of one state to the next one as using OSIRiS construction as initial state and by induction of a single integrase. Cells were induced with either benzoate 100µM for Int5, aTc 200ng/uL for Bxb1 and Arabinose 0.7% for Tp901-1. For characterization every co-transformed bacteria was induced for 16 hours. Each histogram shows fluorescent reporter genes expressed as a result of different induction condition. Each histogram is representative of two different experiments measured by flow cytometry.

**Figure S9.**
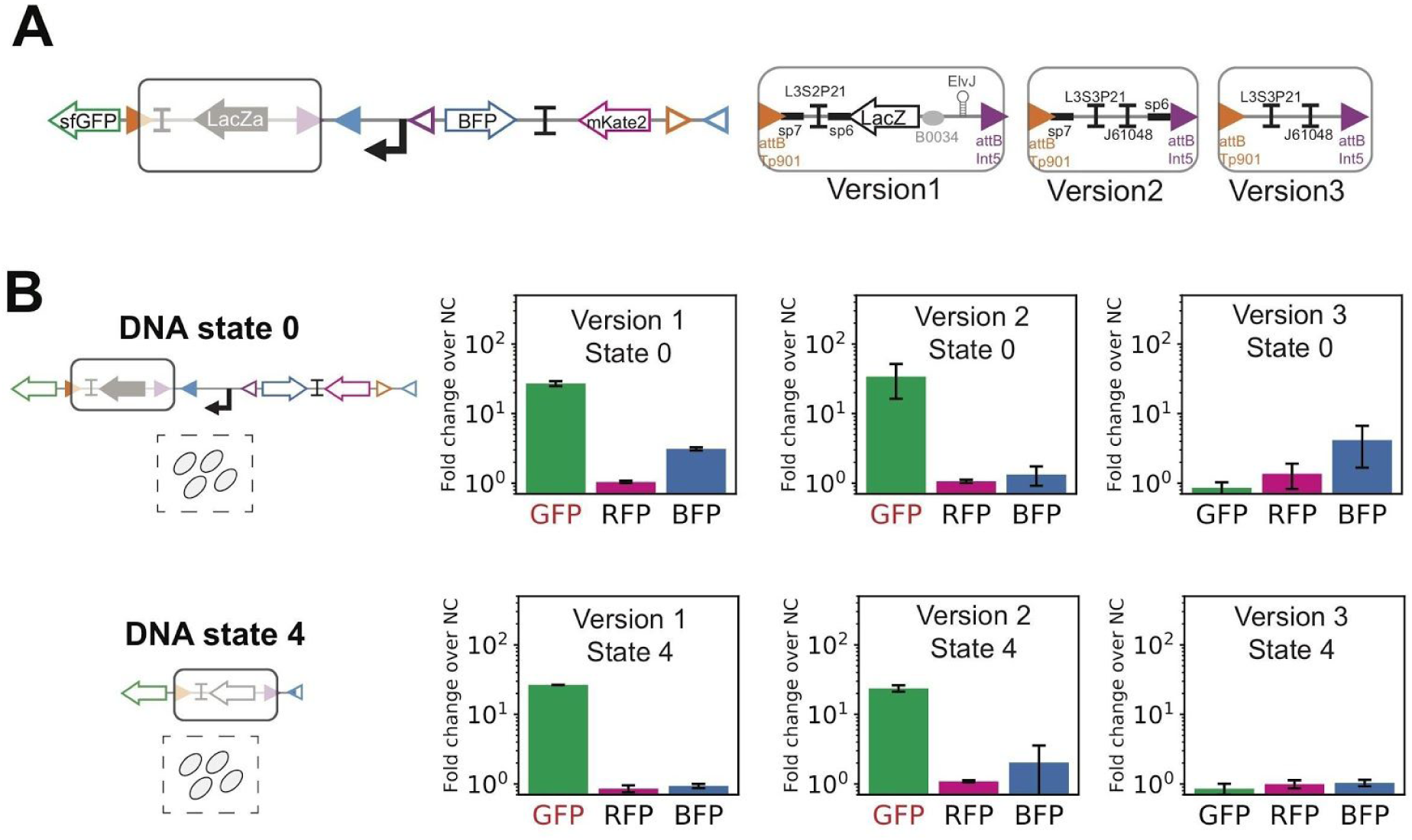
Optimization of 3-input scaffold OSIRiS. **(A)** Generation of two new 3-input OSIRiS designs by modification of the cassette between the attB Tp901 integrase site and the attB Int5 site. For the second and third version, the LacZ gene and 5’UTR were removed and the L3S2P21 terminator was replaced by the two terminators: L3S2P21 an J61048. For version 3, we also removed the two spacers sp7 and sp6. **(B)** Characterization of the initial state and DNA state 4 of the three versions of the 3-input OSIRiS by flow cytometry. We characterized the two selected states via flow cytometry by measuring GFP, RFP, and BFP fluorescence intensities. The bar graph corresponds to the fold change over the negative control (strain without fluorescent protein) for each channel from two experiments with three replicates per experiment. The error bars correspond to the standard deviation between the fold changes obtained in the two separate experiments.

**Figure S10.**
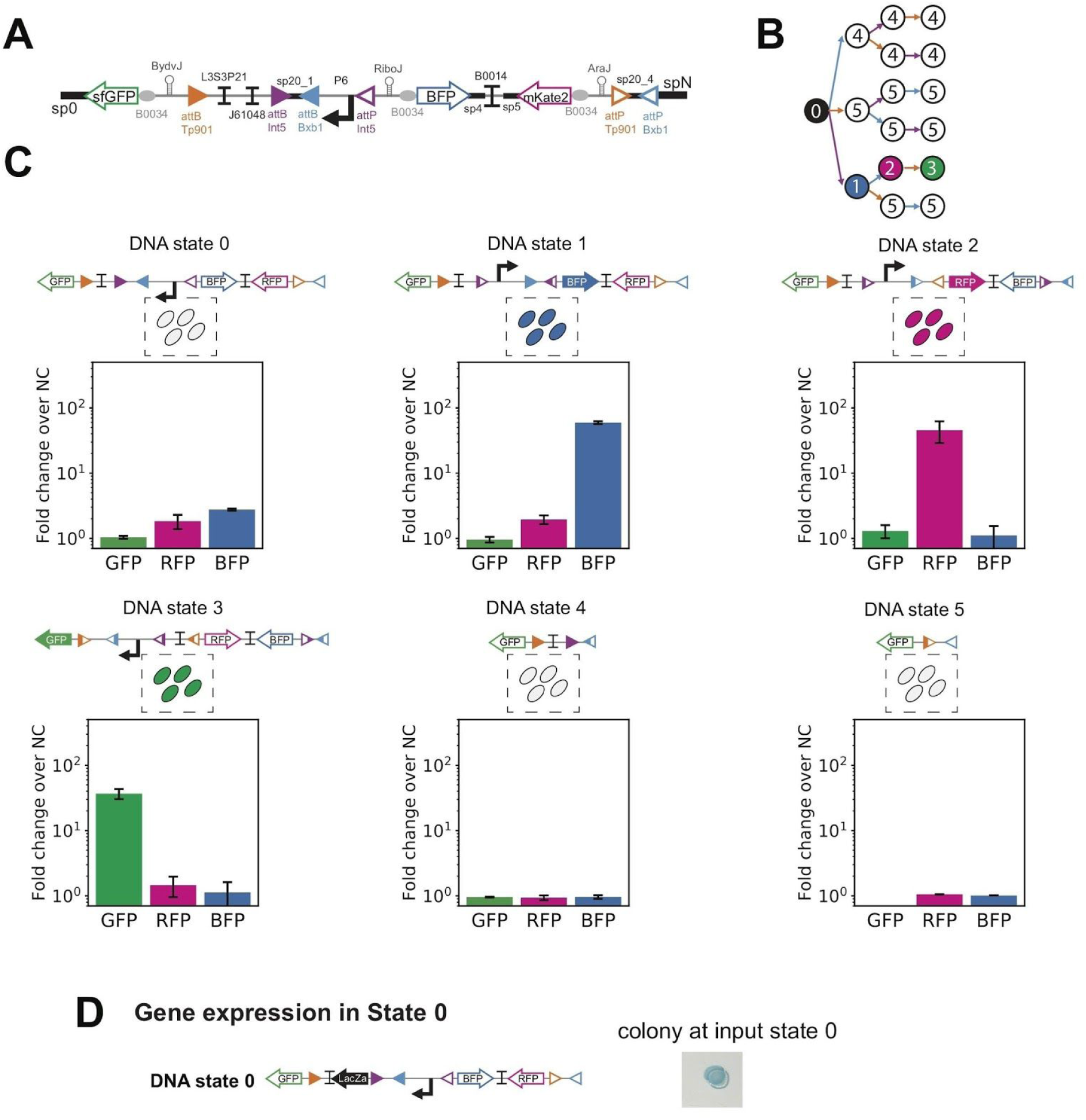
Final design and characterization of 3-input OSIRiS. **(A)** Detailed design for the final 3-input scaffold. In this design, no gene is expressed in the initial input state. **(B)** 3-input lineage tree corresponding to the final 3-input scaffold. The color of each node corresponds to the expected phenotype and the number in the node to the corresponding DNA state of each input state. **(C)** Characterization of the final 3-input OSIRiS by flow-cytometry. We characterized each initial and intermediate recombination states in a flow-cytometry by measurement of GFP, RFP and BFP fluorescence intensities. The bar graph corresponds to the fold change over the negative control (strain without fluorescent protein) for each channel from 3 experiments with 3 replicates per experiment (detailed in Materials and methods). The error bar corresponds to the standard deviation between the fold changes obtained in the 3 separated experiments. **(D)** Additional designed device with *lacZ alpha* gene expression at input state 0, in the 3-input history dependent scaffold. The media was supplemented with X-gal, to obtain blue colonies.

**Figure S11.**
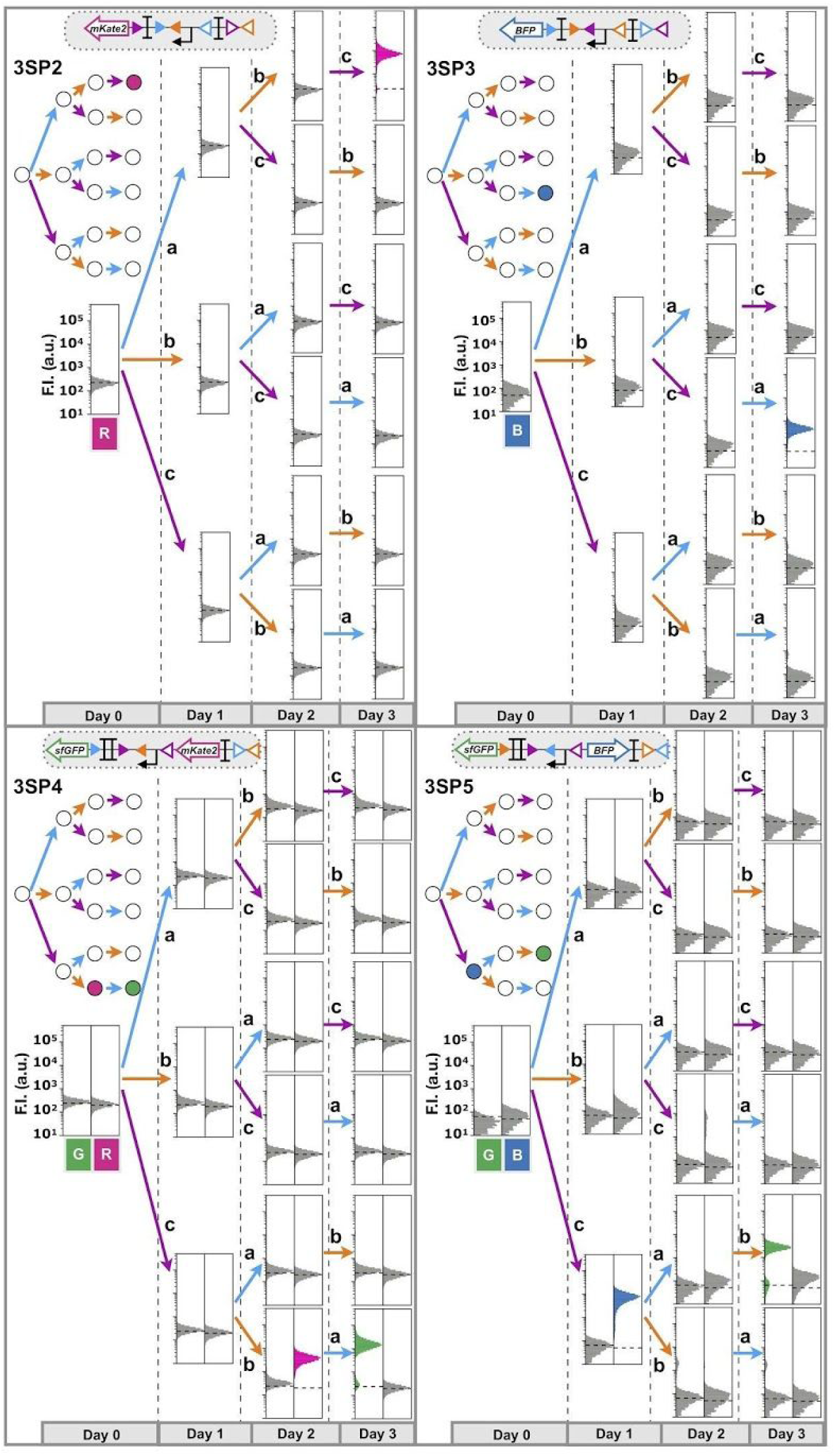
Histograms of 3-input programs single cell characterization. 4 history-dependent single-cell programs were implemented and characterized. We co-transformed each 3-input program with the triple controller plasmid. Bxb1 expression is induced by aTc (input a), Tp901 by arabinose (input b) and Int5 by benzoate (input c). The lineage tree for each program and its corresponding genetic diagram are represented. For characterizing the system, the co-transformed bacteria were induced sequentially for 16 hours each time. Each histogram shows fluorescent reporter genes expressed as a result of different induction conditions corresponding to an input state of the lineage tree. A representative example is depicted here. Fold change measurements can be found in Figure S12.

**Figure S12.**
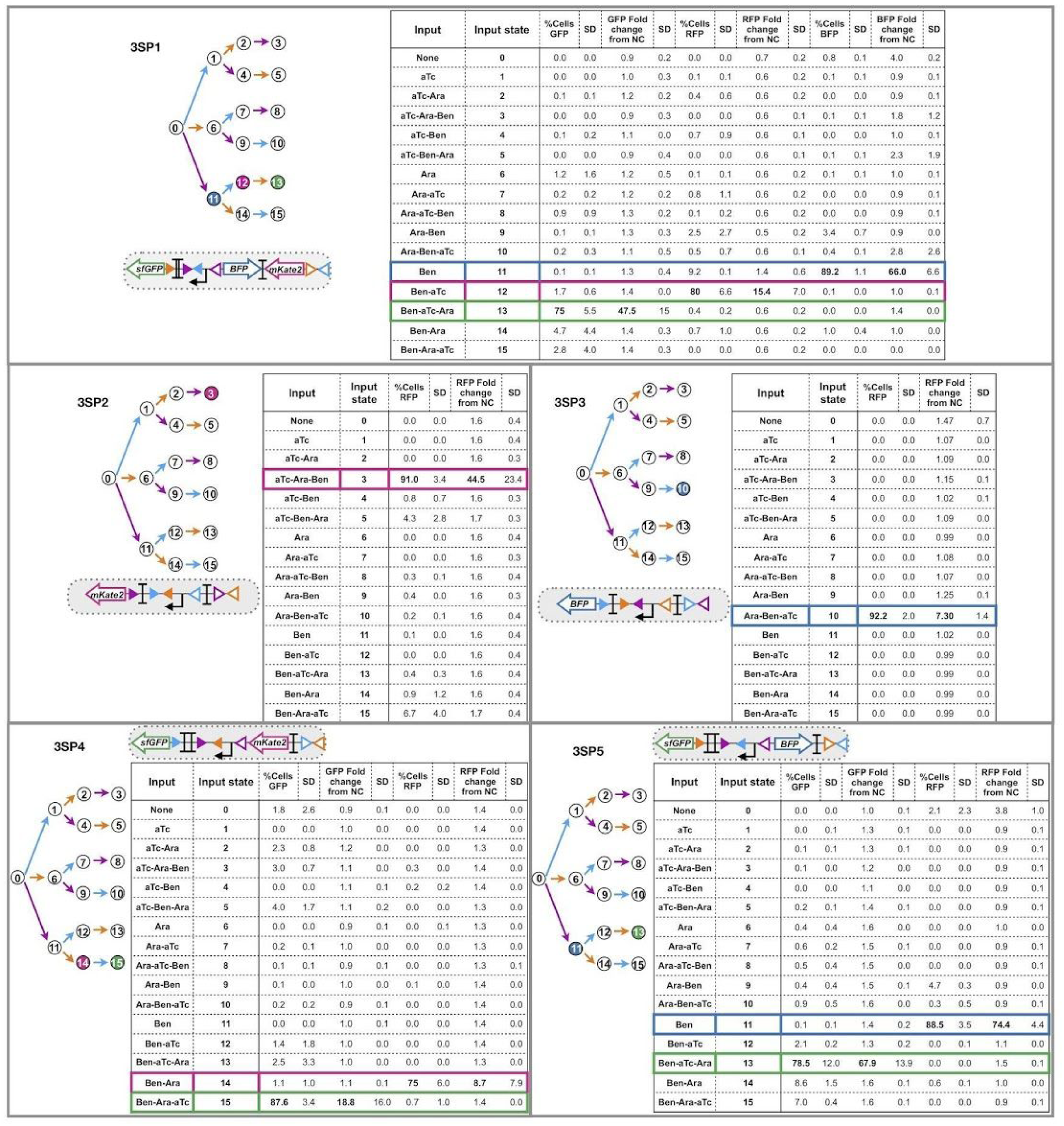
Fluorescence fold change and percentage of cells on different lineage input-states for 3-input single cell programs. The lineage tree for each program and its corresponding genetic diagram are represented. Each table shows % of cell and reporter fluorescence in gates for respective fluorescence channel, as a result of different induction conditions corresponding to an input state of the lineage tree. Values correspond to averages and standard deviations of two different experiments, with tree different colony-forming units analyzed per experiment.

**Figure S13.**
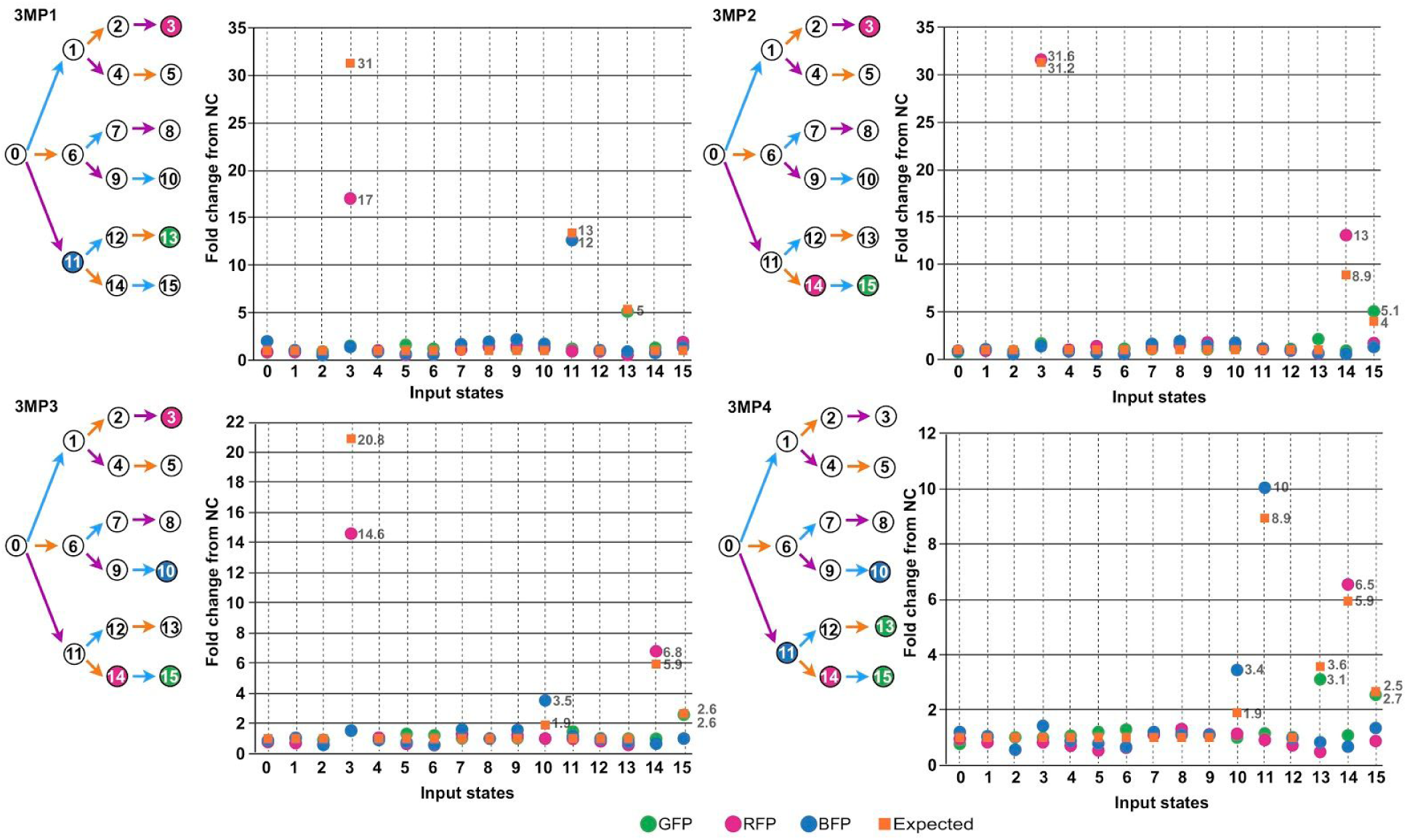
Fold change differences between measured and expected values for 3-input multicellular programs. Plots show the fold change fluorescence over the negative control, resulted for 3-input multicellular programs in every input state. The expected value was obtained dividing single-cell fold change fluorescence values by the number of strains which compose the multicellular system.

**Figure S14.**
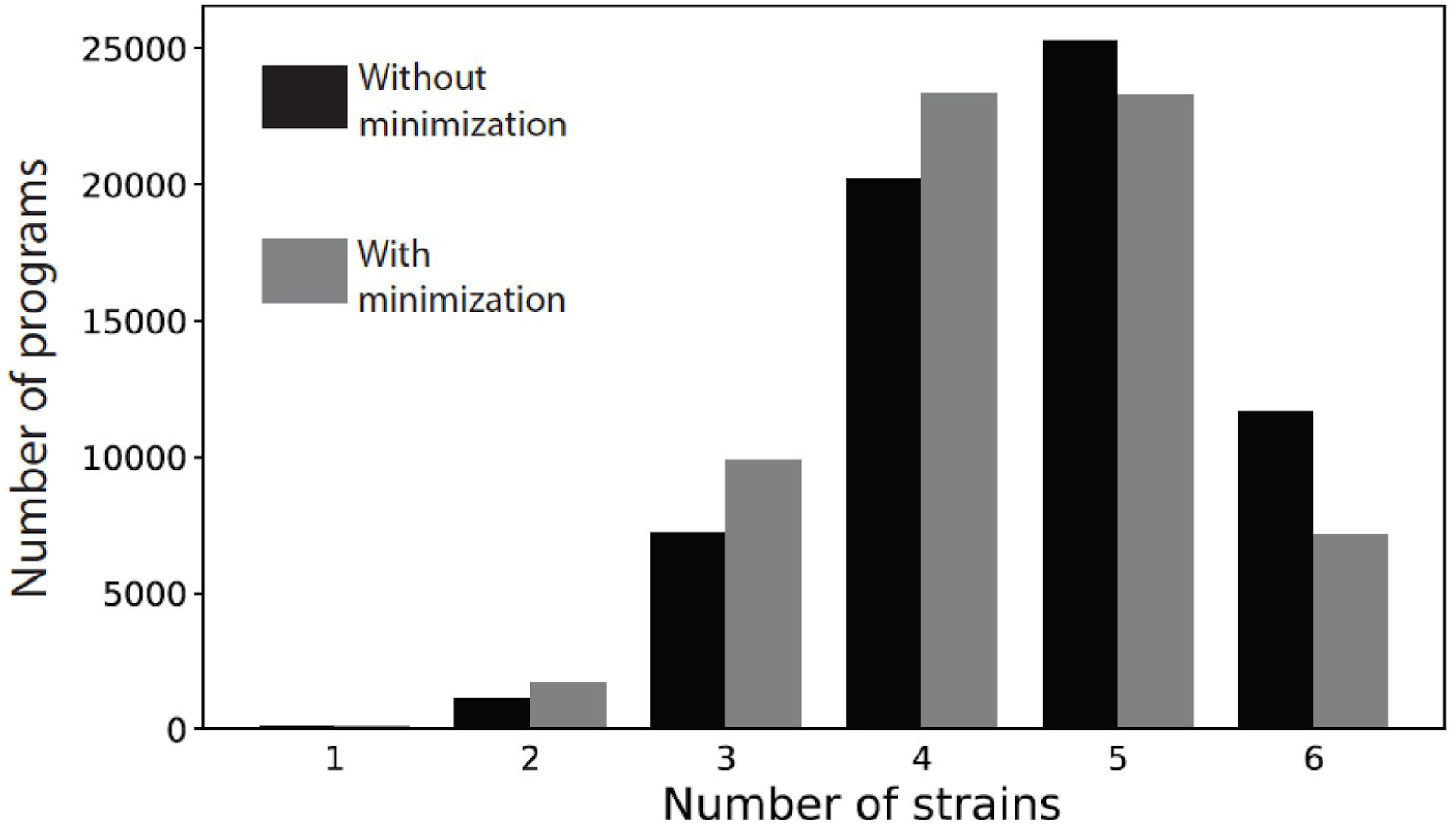
Distribution of the number of strains required for the implementation of all history-dependent programs. Y-axis represents the number of sequential truth table (STT) and X-axis the number of strains required for its implementation. The distribution of strains corresponds to all 3-input 1-output history-dependent programs. Bars show the data without minimization (black bars), and with minimization (grey bars) using Boolean logic devices. Data were obtained using a Python algorithm which generate the designs with the two different strategies for all programs.

**Figure S15.**
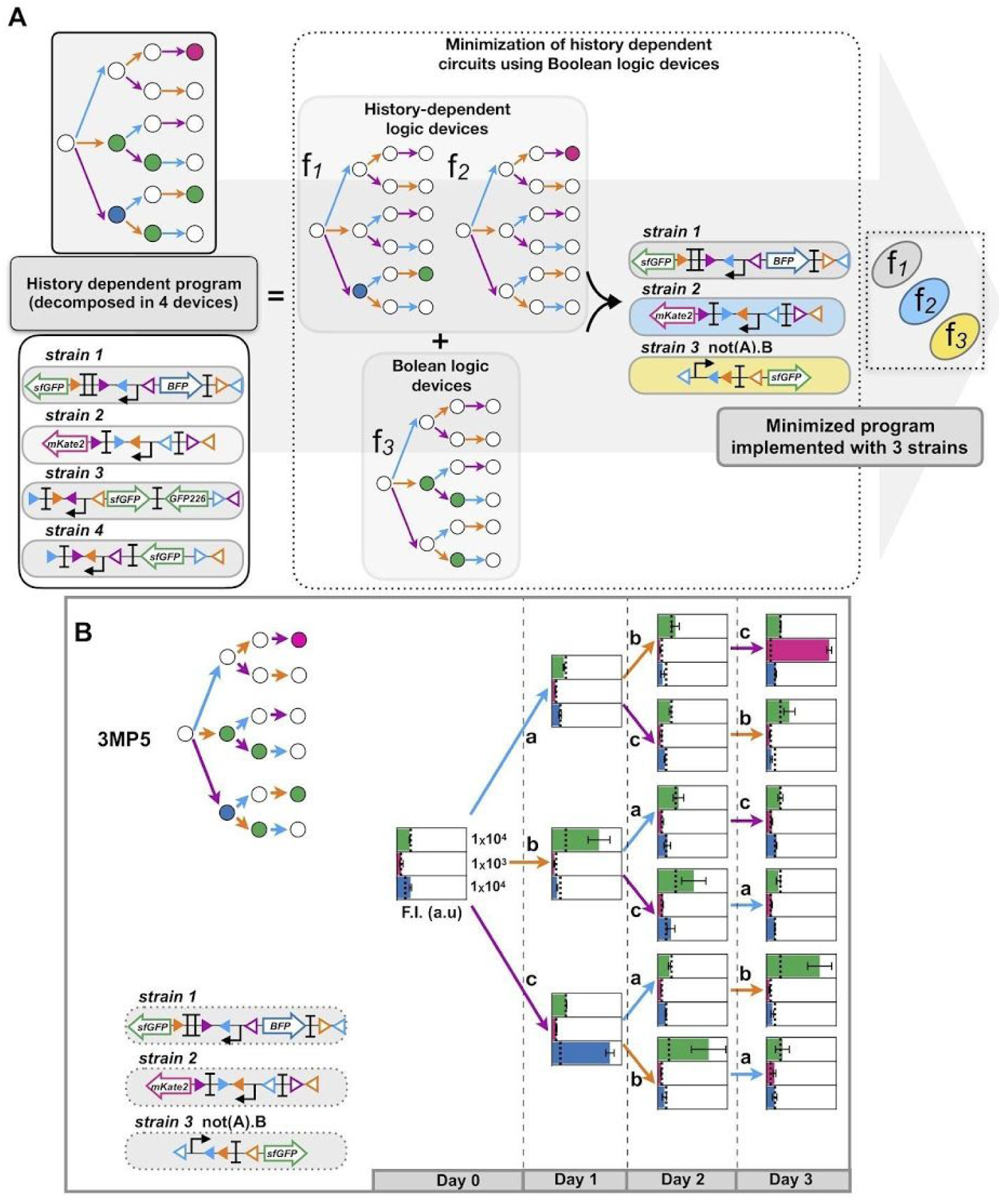
Minimization by simplification of history-dependent programs into Boolean-logic programs. **(A)** Example of a 3-input program (initially decomposed in 4 sub-programs) that can be simplified into 3 sub-programs using Boolean logic devices. Two subprograms correspond to a lineage trees with 2 and 1 ON states in a single lineage. Each sub-program is implemented in differentes strains. The second subprogram corresponds to a lineage tree with 3 ON states in different lineages simplifiable into a Boolean-logic function (not(A).B), implemented in one strain. Using this minimization scheme, we minimized the required number of strains from 4 to 3. **(B)** Characterization of the 3-input multicellular program simplified on A. The sub-programs were implemented in 3 different strains. The inputs are represented by letters, a for Integrase Bxb1, b for integrase Tp901-1 and c for Int5. The strains were mixed in similar proportions and grown consecutively during 16 hours with each inductor. Each graph corresponds to a different induction condition corresponding to an input state of the lineage tree. The bar graph corresponds to the fluorescence intensity (F.I) in arbitrary units (a.u) for each fluorescent chanel (GFP, RFP and BFP). The error bars correspond to the standard deviation for three biological replicates. The dotted line indicates the negative control autofluorescence.

**Figure S16.**
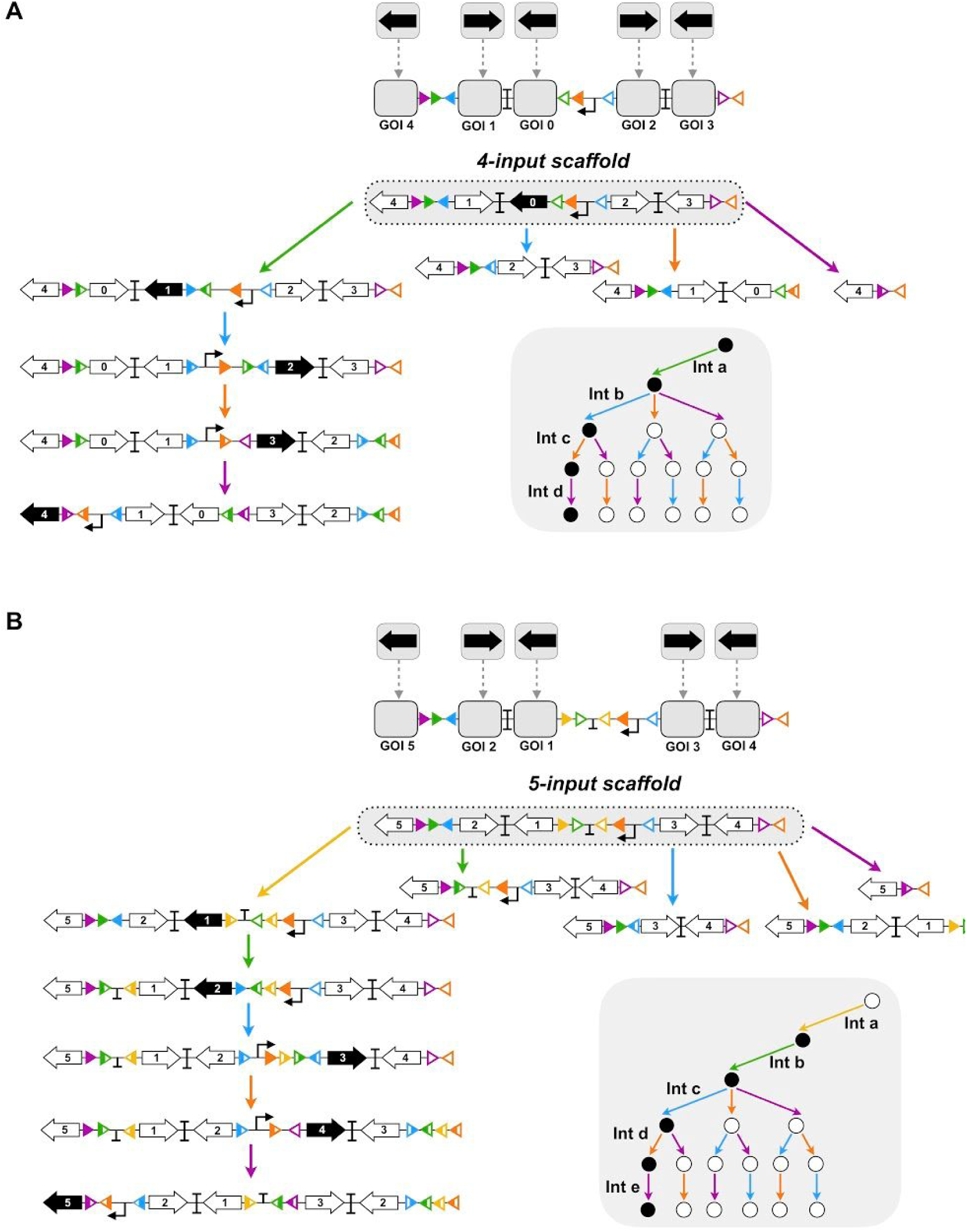
4 and 5 input scaffold and states diagrams. 4-input **(A)** and 5-input **(B)** scaffolds. Integrase sites are positioned in each scaffold to permit expression of an output gene in the corresponding lineage. Therefore, for each state of the lineage a different gene is expressed. On the right panel A, gene 0 is expressed only when no input is present. If input a is present first, gene 1 is expressed, but if input b, c or d are present first, none gene is expressed (nor will be expressed) as the promoter is excised. If input b follows input a, gene 2 is expressed and so on. The 5-input scaffold allows expression of a different GOI in each state except in the state 0 (with no input). An additional strain is needed if gene expression is required in this state. Here only intermediate states for the scaffold lineage are represented. For other lineages only the first intermediate state is represented.

**Figure S17.**
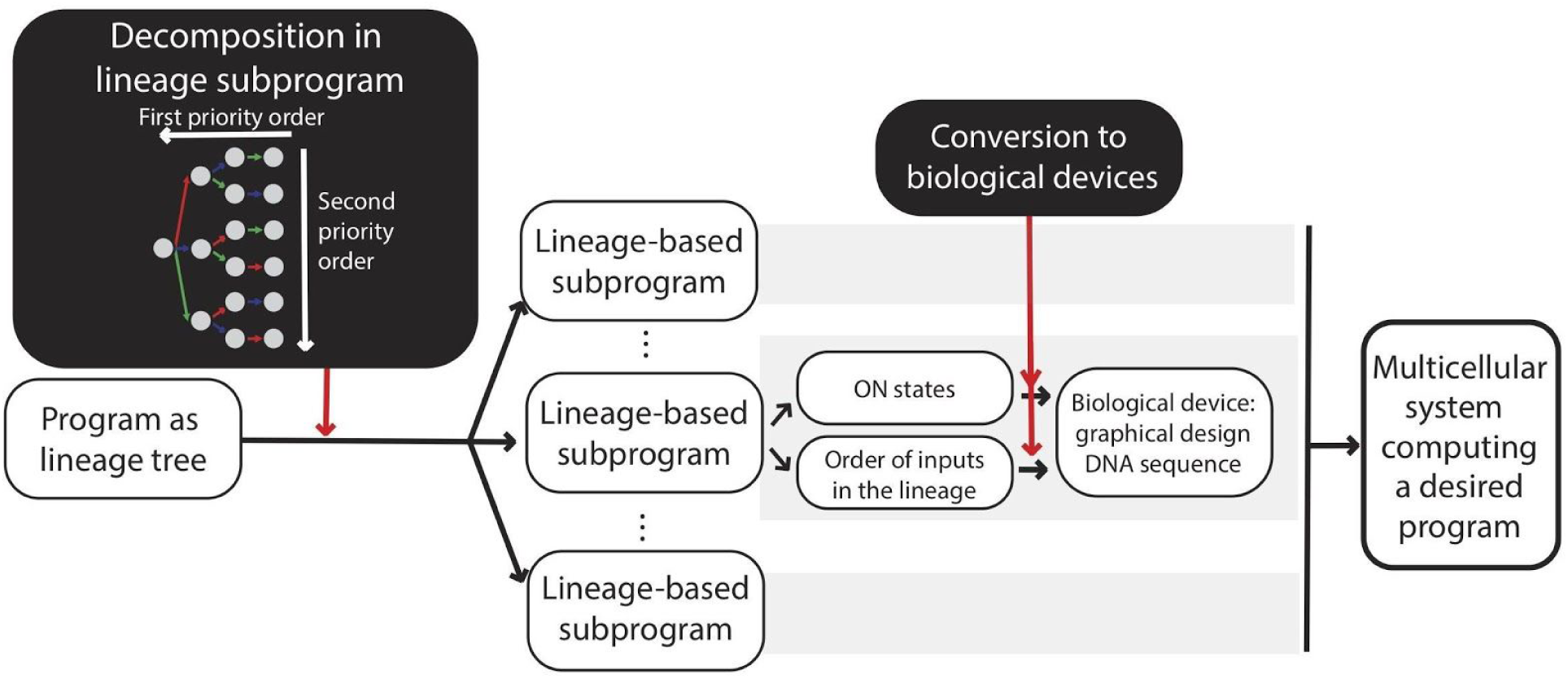
Automated design of history dependent programs using CALIN web-interface. The Python program takes as input a history-dependent program written as a lineage tree. This program is decomposed into sub-programs, and the decomposition is performed by preferentially extracting subprograms with ON state at the extremity of the tree (corresponding to state with the highest number of inputs present). For each subprogram, the algorithm identifies the identity of ON states and the order of the inputs in the lineage. Based on this information, the biological design is obtained and the graphical design for integrase and history-dependent DNA devices with its corresponding DNA sequence are given. The full program composition is implemented by combining the population of strains with different subprograms.

**Figure S18.**
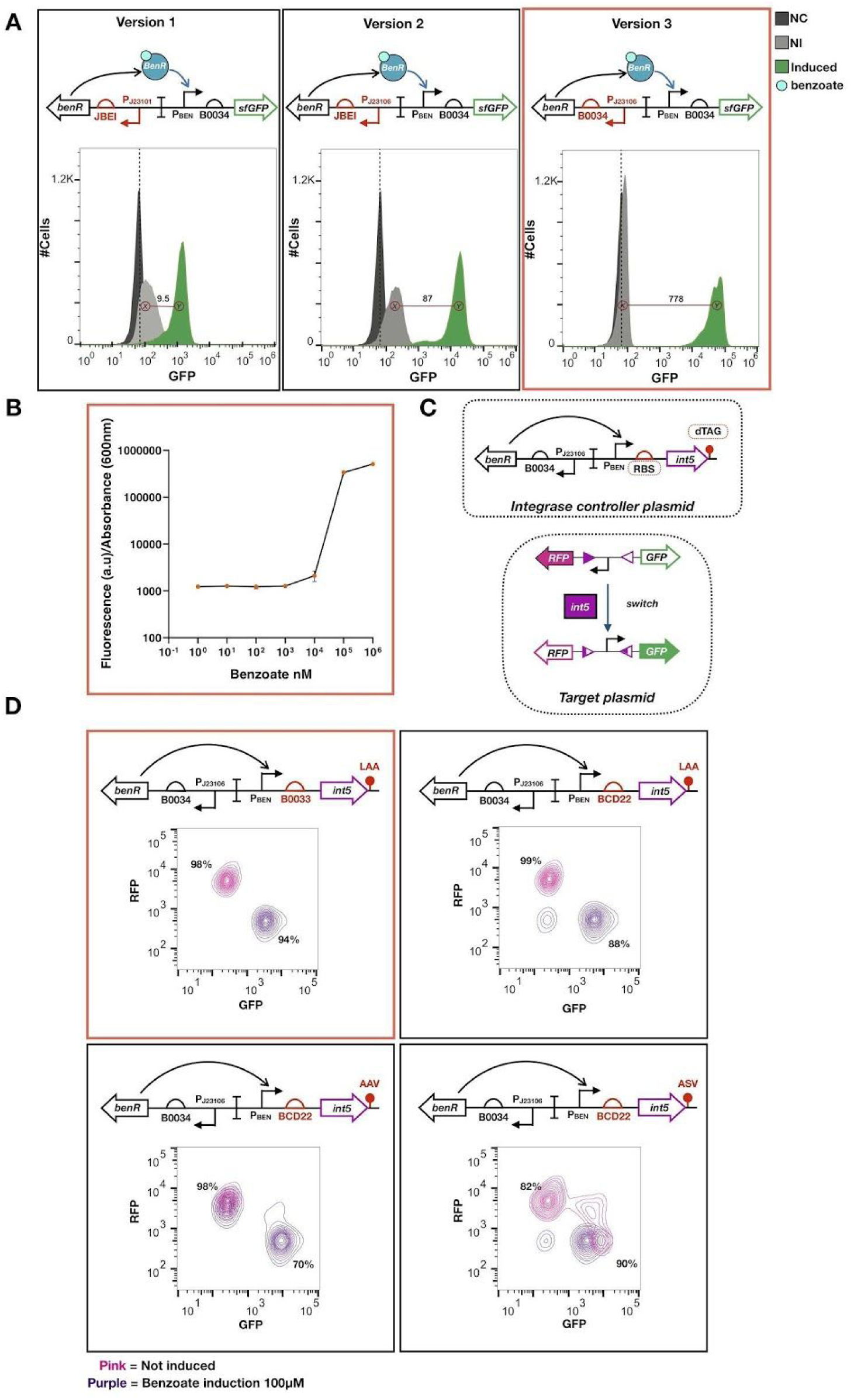
Optimization of BenR and Integrase 5 expression. Three genetic versions of benzoate sensor BenR were characterized (A). The constructions were designed with different combinations of promoters and RBS to tune *benR* gene transcription, and induce GFP fluorescence in presence of benzoate. These versions were characterized using 10 μM of benzoate for version 1 and 2 and 100 μM for version 3 as inductor for 16h at 37°C. Higher concentration affected the growth of cell in versions 1 and 2. The fold change between Non-induced cells population (NI, *X* population) and Induced cells population (*Y*) was calculated, for each. Titration curve response of sensor version 3 to benzoate (B). Cell harboring the version 3 of the sensor were induced with different benzoate concentration for 16h. The fluorescence was measured by plate reader. A maximum fluorescence was observed after 100 μM of benzoate induction. Scheme of Integrase 5 gene expression optimization and target plasmid to test switching efficiency (C). Characterization of different constructions for *int5* gene expression (D). Different RBS and degradation tags were tested. Plots show flow cytometry measurements of fluorescence. Cells not induced (pink) are expressing RFP fluorescence (state 0). After induction with benzoate 100 μM (purple), expression of GFP is observed (state 1). The percentage of cells in each state are showed. Red square indicates the construction selected to use for integrase triple controller plasmid.

**Table Supplementary1.**

**Table Supplementary 1**
| Program name | Minimun Fold Change |  |  |
| --- | --- | --- | --- |
|  | GFP | RFP | BFP |
| **2SP1** | 12 | - | - |
| **2SP2** | 8 | - | - |
| **2SP3** | 7 | - | - |
| **2SP4** | 10 | - | - |
| **2SP5** | - | 26 | - |
| **2SP6** | 37 | - | - |
| **2SP7** | 37 | 25 | - |
| **2SP8** | 39 | - | 130 |
| **2SP9** | 23 | 103 | 199 |
| **3SP1** | 33 | 10 | 16 |
| **3SP2** | - | 26 | - |
| **3SP3** | - | - | 7 |
| **3SP4** | 18 | 6 | - |
| **3SP5** | 42 | - | 20 |

The minimum fold change corresponds to the fold change between the ON state and the maximum OFF state

